# Gas-vacuolate *Microcystis* evolves cyanophage resistance under low nitrogen conditions

**DOI:** 10.64898/2026.07.12.738045

**Authors:** Isaac Meza-Padilla, Jozef I. Nissimov

## Abstract

Cyanophages can influence the dynamics of toxic cyanobacterial blooms. However, cyanobacteria can become resistant to viruses through natural selection processes. Here, we investigate the acquisition of virus resistance in a toxic, freshwater, gas-vacuolate, bloom-forming cyanobacterium, *Microcystis aeruginosa*, under different nutrient concentrations. We find that gas-vacuolate *M. aeruginosa* subpopulations acquire virus resistance in low nitrogen cultures regardless of their phosphorus concentration, whereas non-vacuolate subpopulations do not. After resequencing susceptible and resistant *M. aeruginosa* variants, we identify a mutation in the transmembrane domain of a nitrogen-related transporter as the most likely genetic cause of the resistance. Infection experiments further reveal a larger viral burst size and higher phycocyanin content in gas-vacuolate cells compared to non-vacuolate ones. Based on these experimental results, we propose an ecological model in which lower nitrogen concentrations, higher light intensities and increased virus-host contact rates facilitate the evolution of virus resistance in upper lake layers during *Microcystis*-dominated blooms.

## Introduction

In 1996, an outbreak of acute liver failure took place at a hemodialysis clinic^1,2^. During the outbreak, 116 patients experienced symptoms such as muscle weakness, nausea, visual disturbances and vomiting after routine haemodialysis treatment^3^. 100 patients developed acute liver failure and 76 of them died^4^. Liver tissue and sera analyses revealed that 52 of the victims had been exposed to lethal levels of the cyanobacterial hepatotoxin microcystin^5^. These events occurred after using water from a local reservoir that was experiencing harmful algal blooms dominated by toxic cyanobacteria, including *Microcystis*^2,4^.

The above is but one example of the devastating consequences of cyanobacterial harmful algal blooms (cyanoHABs) in freshwater resources. Other deleterious societal impacts of toxic cyanoHABs include the deaths of cattle^6^, dogs^7,8^, and commercially important fish species^9,10^. Many bloom-forming cyanobacteria produce toxins that affect the nervous system or induce liver damage^11,12^. Beyond their toxigenic effects, cyanoHABs also cause noxious hypoxic events in freshwater bodies due to elevated respiration rates during bloom decomposition, which lead to increased mortality and biodiversity loss among aquatic organisms^13,14^. Furthermore, cyanoHABs are becoming increasingly common worldwide in stratified and non-stratified lakes due to several factors, including anthropogenically-driven eutrophication and the resulting increases in nitrogen and phosphorus concentrations in lakes^15–17^.

Several factors cause a negatively correlated nutrients-light gradient in stratified lakes. Briefly, as depth increases, the concentration of nutrients increases and the light intensity decreases^18^. This phenomenon in turn directly impacts cyanobacteria, including *Microcystis*, which need to optimize their vertical position in the water column in order to acquire sufficient light and nutrients for growth^19^. One of the most remarkable adaptations of cyanobacteria for buoyancy regulation is the gas vacuole, an aggregation of hollow proteinaceous structures called gas vesicles^20^. In brief, gas vacuoles confer buoyancy to cyanobacterial cells, whereas non-vacuolate morphotypes sink^21^.

Lytic cyanophages (i.e., viruses that infect and kill cyanobacteria) can influence the dynamics of their hosts and have been linked to the termination of cyanoHABs^22–24^. Viruses have even been considered potential biological agents to control cyanoHABs^25,26^. The high host density within cyanoHABs creates a favourable scenario for lytic viral infections, particularly in standing bodies of water such as lakes. However, cyanobacteria can become resistant to viruses through natural selection processes^27–30^, thereby challenging our understanding of how viral infections influence cyanoHAB dynamics and bloom termination.

The lytic, linear double-stranded DNA Ma-LMM01 virus infecting the toxic, bloom-forming *M. aeruginosa* NIES-298 is the most comprehensively characterized freshwater cyanophage-host system to date^31–33^. Conveniently, the *M. aeruginosa* NIES-298 host strain is composed of gas-vacuolate and non-vacuolate cells (see Results). In this study, we set up a factorial culture matrix comprised of different nitrogen and phosphorus concentrations in order to compare Ma-LMM01/NIES-298 infection dynamics under varied concentrations of nutrients. While doing so, two NIES-298 variants cultured under low nitrogen conditions became resistant to Ma-LMM01, thus providing a unique opportunity to experimentally investigate the acquisition of virus resistance in freshwater cyanobacteria within the context of eutrophication, buoyancy morphotypes, and toxic cyanoHABs.

## Results

### Nutrients infection experiment

After a year of semi-continuous culturing of *M. aeruginosa* NIES-298 under different nitrogen and phosphorus concentrations, namely, high N & high P (hN-hP), low N & high P (lN-hP), high N & low P (hN-lP), and low N & low P (lN-lP), we infected NIES-298 cultures with the cyanophage Ma-LMM01 to compare their infection dynamics under the aforementioned nutrient conditions. During this infection experiment, it came to our attention that the cyanobacterial cultures under low N conditions did not undergo complete lysis (**Fig. 1**a-d). Statistical analyses applied to the final day of the experiment (i.e., 3 days post infection; dpi) indicated that there was a significant interaction between N and P regarding the cell density (log_10_) of infected cultures (two-way ANOVA, *F*_1,8_ = 22.48, *P* = 0.001). Tukey’s HSD confirmed that the cell density of infected low N cultures was significantly higher compared to that of cells grown in infected high N conditions (low N vs high N groups, *P* < 1 × 10^−7^ in all cases). In contrast, no difference was detected between infected high N cultures (hN-hP vs hN-lP, *P* = 0.696), which had lysed by the end of the experiment. The interaction between N and P was also present with respect to the maximum quantum yield of photosystem II (Fv/Fm), which is an indicator of photosynthetic efficiency, during the last day of infection (*F*_1,8_ = 275.5, *P* = 1.75 × 10^−7^). As with cell density, a Tukey’s post-hoc test identified significant Fv/Fm differences between low N and high N cultures (*P* ≤ 1 × 10^−7^ in all cases), but not between high N groups (hN-hP vs hN-lP, *P* = 0.899). As expected, in the uninfected control cultures the cell density increased, and the photosynthetic efficiency of the cyanobacterial cells remained constant.

**Fig. 1.**
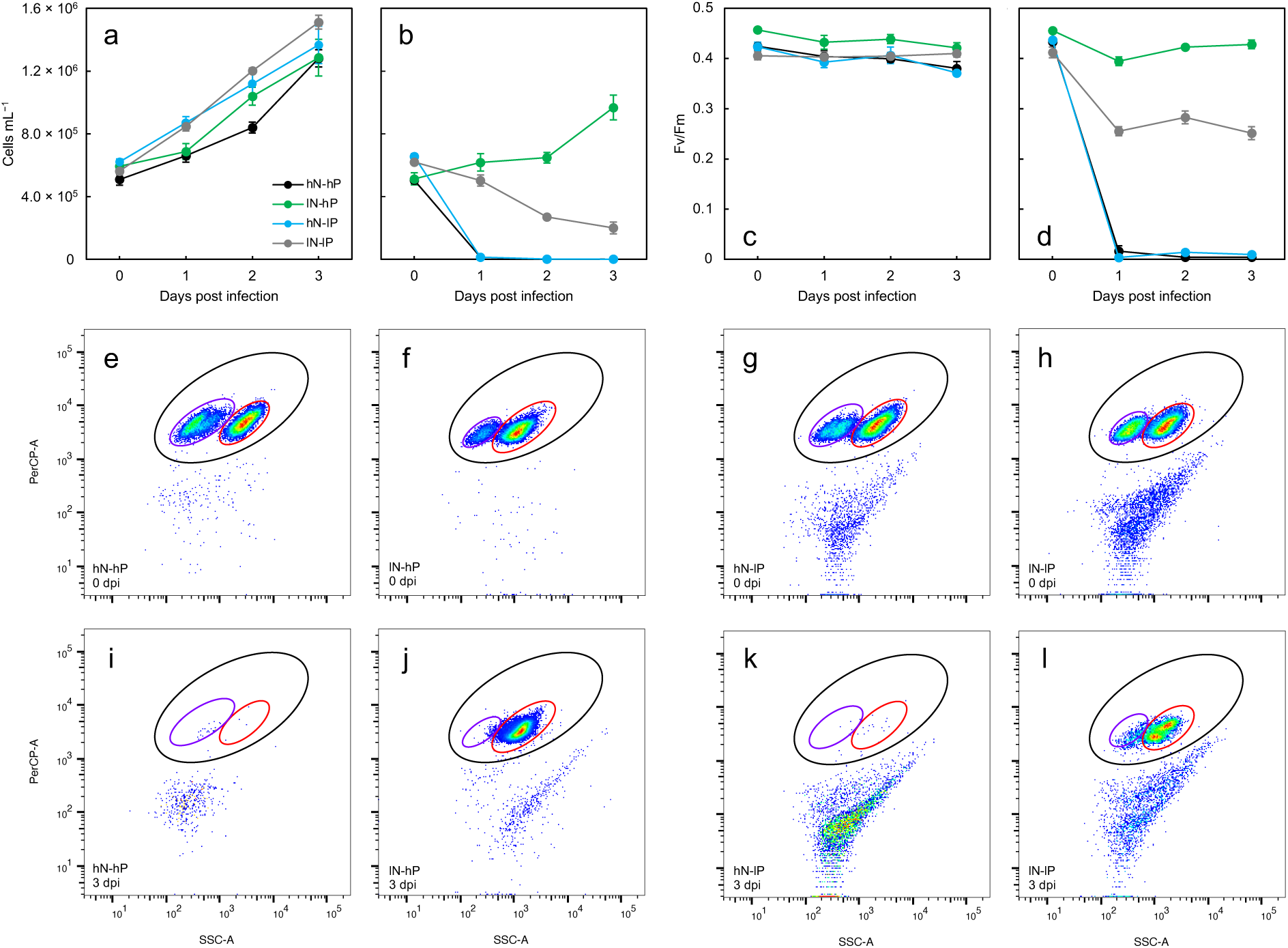
Nutrients infection experiment. (a) Cells mL^−1^ in uninfected *Microcystis aeruginosa* NIES-298 cultures over the course of the experiment, and (b) cells mL^−1^ in cultures infected with Ma-LMM01. (c) Maximum quantum yield of photosystem II (Fv/Fm) in uninfected cultures, and (d) Fv/Fm in infected cultures. Black, green, light blue, and gray dots indicate the mean (*n* = 3) of high N & high P (hN-hP; i.e., standard BG-11 media), low N & high P (lN-hP), high N & low P (hN-lP), and low N & low P (lN-lP) groups, respectively. Error bars denote ± standard deviations. (e-h) And (i-l) display live cytograms of infected NIES-298 cultures 0 days post infection (dpi) and 3 dpi, respectively. hN-hP are shown in (e,i), lN-hP in (f,j), hN-lP in (g,k), and lN-lP in (h,l). Black gates enclose the entire NIES-298 population, purple gates the NIES-298 subpopulation with the weaker side scatter signal, and red gates the subpopulation with the stronger side scatter. The density of the events is displayed using a blue-red gradient, blue indicating the lowest density and red the highest. PerCP-A, red fluorescence (log scale); SSC-A, side scatter (log scale).

Visual examination of flow cytometry (FC) plots 0 dpi readily revealed two NIES-298 subpopulations in all treatments (**Fig. 1**e-l). Both subpopulations had the same red fluorescence (i.e., the same chlorophyll concentration) but differed in their side scatter (a measure of intracellular complexity) properties. Interestingly, we observed that most of the cyanobacterial cells resistant to Ma-LMM01 in low N cultures were part of the subpopulation with the stronger side scatter signal. Statistical analyses showed that the percentage change in cell density from 0 to 3 dpi differed significantly between the two subpopulations in both lN-hP (paired *t*-test, *t*_2_ = 20.696, *P* = 0.002) and lN-lP (*t*_2_ = 8.367, *P* = 0.014) treatments. By the end of the experiment, the percentage change in cell density was 210.25 ± 16.68% (mean ± s.d.) in the stronger side scatter NIES-298 subpopulation in infected lN-hP cultures, compared to 6.59 ± 0.37% in the weaker side scatter subpopulation. In lN-lP cultures, the corresponding percentage changes were 48.91 ± 9.75% and 4.36 ± 0.53%, respectively.

### Cell sorting and morphological observations

In order to investigate the morphological differences between the two NIES-298 subpopulations, we started by separating them into different cultures. Subpopulations susceptible to Ma-LMM01 were isolated via cell sorting, while resistant subpopulations were isolated by infecting low N NIES-298 cultures with the virus and culturing the surviving cells. This was done in order to ensure that we were isolating only resistant cells and no susceptible ones. Once the two subpopulations were separated and re-cultured, we analyzed the resulting cultures again through FC and observed the cells under the microscope. Only one subpopulation was observed in all cases (**Fig. 2**), indicating that the separation and isolation of subpopulations was successful. Brightfield microscopy revealed that the NIES-298 cells that presented the stronger FC side scatter signal (more intracellularly complex) contained gas vacuoles (**Fig. 2**). In contrast, the NIES-298 cells that displayed the weaker FC side scatter signature (less intracellularly complex) did not contain gas vacuoles. These morphological observations were also supported by the overall appearance of the cultures themselves in their static non-shaken state: cyanobacterial cells in gas-vacuolate cultures floated and were distributed throughout the flasks, whereas those in non-vacuolate cultures sank to the bottom (**Fig. 2**). Taken together, these results indicated that resistance against Ma-LMM01 emerged in gas-vacuolate cells cultivated in low N conditions.

**Fig. 2.**
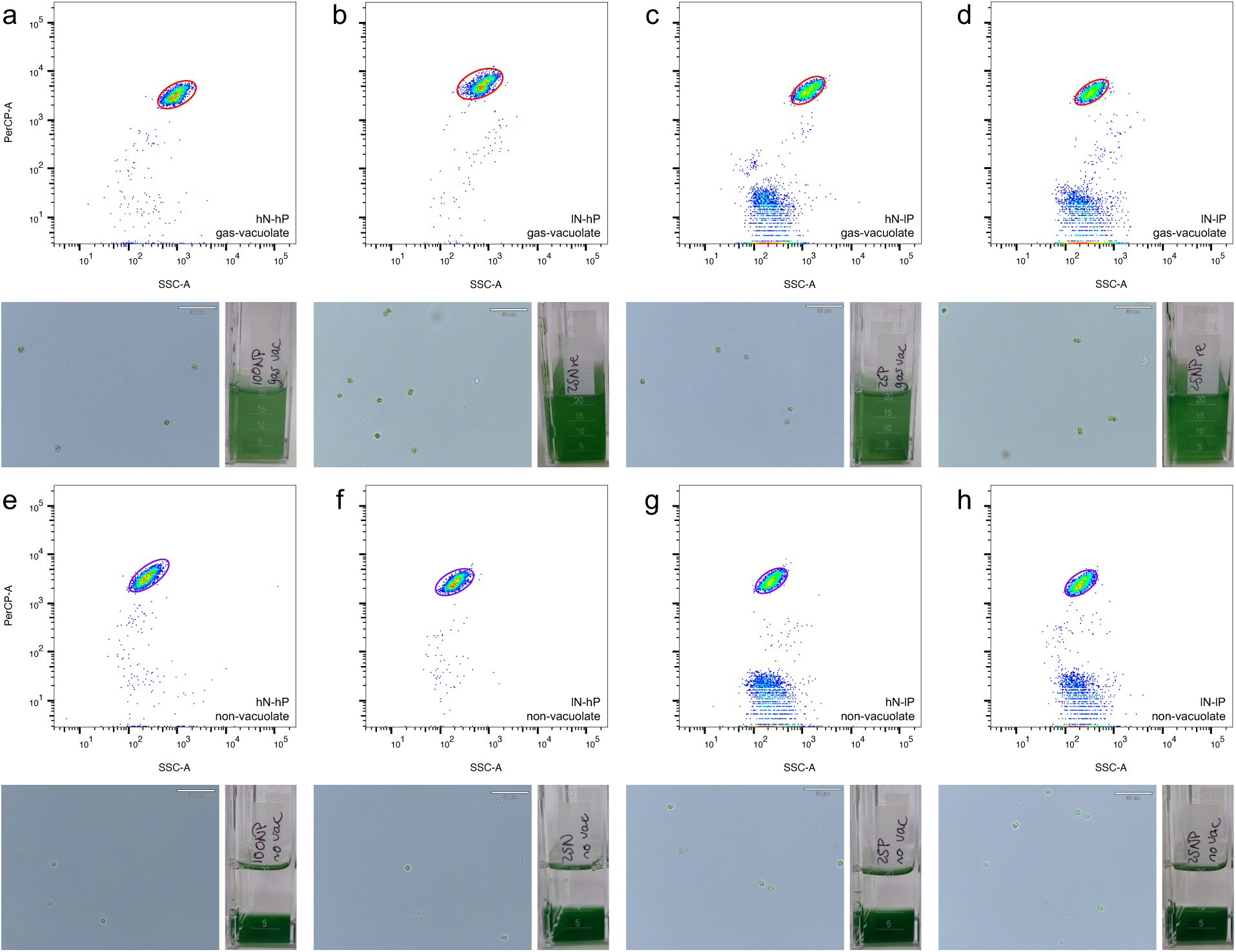
*Microcystis aeruginosa* NIES-298 cultures sorted into distinct subpopulations. (a-d) Show hN-hP, lN-hP, hN-lP, and hN-hP cultures, respectively, containing gas-vacuolate NIES-298 cells. (e-h) Display hN-hP, lN-hP, hN-lP, and hN-hP cultures, respectively, containing non-vacuolate NIES-298 cells. The panels show live cytograms, brightfield micrographs, and photographed flasks of the aforementioned cultures. The cytograms are plotted using red florescence (PerCP-A) vs side scatter (SSC-A) in logarithmic scale. The density of the events is displayed using a blue-red gradient, blue indicating the lowest density and red the highest.

### Mutation analyses

We re-sequenced the genomes of the eight separated NIES-298 subpopulations (i.e., gas-vacuolate and non-vacuolate hN-hP, lN-hP, hN-lP, and lN-lP cultures) to investigate genomic changes associated with resistance against Ma-LMM01. Prior to sequencing, the separated NIES-298 subpopulations were infected with Ma-LMM01 in order to verify their susceptibility or resistance. This confirmed our previous findings that gas-vacuolate cells from low N cultures were resistant to Ma-LMM01, while the remaining six subpopulations were susceptible to the virus (**Fig. S1**).

We reasoned that the mutations associated with resistance would be those shared between resistant NIES-298 subpopulations and absent in susceptible ones. Three genetic variants met these criteria (**Table 1**). The first one was a cytosine deletion in an intergenic homopolymeric C tract 199 bp upstream of a PEP-CTERM sorting domain-containing protein and 259 bp upstream of a Uma2 family endonuclease. The second was a nonsynonymous SNP in the amidophosphoribosyltransferase gene (*purF*; RefSeq accession number: WP_103113045.1) of NIES-298, resulting in a leucine-to-valine substitution at position 71. The third was a nonsynonymous SNP that resulted in a glycine-to-valine change at position 554 in a transmembrane protein exporter (annotated in RefSeq as a peptidase domain-containing ABC transporter; WP_103113363.1).

**Table 1.** Mutations shared between Ma-LMM01-resistant *Microcystis aeruginosa* NIES-298 subpopulations (i.e., gas-vacuolate cells from low N cultures) and absent in susceptible ones.

| Position<sup>a</sup> | Mutation<sup>b</sup> | Description | Gene | Annotation | Frequency<sup>c</sup> |
| --- | --- | --- | --- | --- | --- |
| 3,210,317 | (C)<sub>12</sub>→<sub>11</sub> | Intergenic (-199/-259) | ACNQK5_RS15405 ← / → ACNQK5_RS15410 | PEP-CTERM sorting domain-containing protein/Uma2 family endonuclease | 94.0-96.0, 152-178 |
| 4,213,818 | G→C | L71V (CTT→GTT) | *purF* ← | Amidophosphoribosyltransferase | 99.8-100, 458-485 |
| 3,802,561 | G→T | G554V (GGT→GTT) | ACNQK5_RS18290 → | Peptidase domain-containing ABC transporter | 99.5-99.8, 390-408 |

<sup>a</sup>All mutations occur in the bacterial chromosome.
<sup>b</sup>Nucleotide substitutions (‘Mutation’ column) are reported relative to the reference genome sequence, and codon changes (‘Description’ column) are shown in coding-strand orientation.
<sup>c</sup>Values before the comma indicate the percentage of reads supporting the mutation, and values after the comma indicate the total read count at that position, as reported by *breseq*.

During these analyses, we also detected mutations that were present exclusively in gas-vacuolate NIES-298 subpopulations. All gas-vacuolate NIES-298 had mutations either in the intergenic region between the gas vesicle protein-coding genes *gvpC* and *gvpA*, within a *gvpA* gene, or both (**Table S1**). These variants were absent in non-vacuolate cells. The only gas vacuole-related mutation observed in a non-vacuolate subpopulation (hN-lP) was a SNP in a *gvpA* gene that converted the canonical start codon ATG to ATA, likely impairing translation initiation.

### Structural analyses and adsorption assays

We proceeded to investigate the structural effects of the mutation that occurred in the protein exporter of the NIES-298 subpopulations resistant to Ma-LMM01. This nitrogen-related transporter was predicted to contain a transmembrane domain with six transmembrane α-helices. The mutation occurred within the transmembrane domain of the protein exporter, in an intracellular amino acid located right before the start of the third (counting from N-terminus to C-terminus) transmembrane α-helix (**Fig. 3**a,b). The mutation changed glycine 554, a small and flexible residue, to valine, a bulky and hydrophobic one. Not surprisingly, this caused a major change in the secondary structure: the original bend (coil) formed by the lysine was replaced by a helix (**Fig. 3**c). Missense3D further supported this result, classifying the mutation as damaging after replacing a glycine originally located in a bend curvature. In addition, the lysine-to-valine substitution drastically affected the local hydrophobicity potential, changing the molecular surface from hydrophilic to lipophilic (**Fig. 3**d). Both effects were observed before and after energy minimization.

**Fig. 3.**
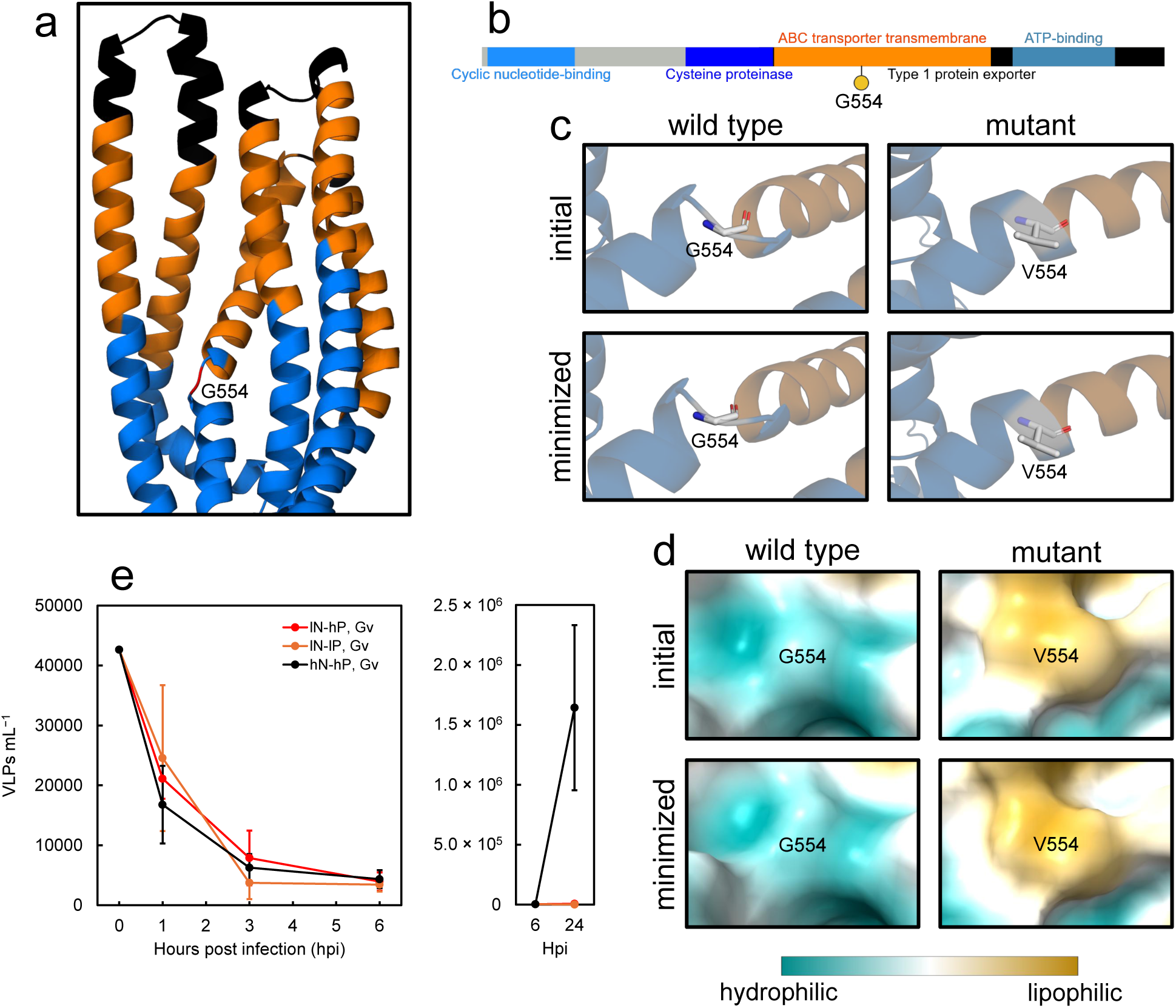
Structural effects of the nitrogen-related transporter mutation that occurred in *Microcystis aeruginosa* NIES-298 subpopulations resistant to Ma-LMM0, and adsorption assays. (a) Structural model of NIES-298 protein exporter. The intracellular region is colored in blue, the transmembrane α-helices in orange, and the extracellular region in black. Glycine 554 is colored in red and labeled. (b) Schematic of the domains present in the transporter. Glycine 554 is highlighted with a yellow circle. (c) Changes in secondary structure due to the G554V mutation, before and after energy minimization. Atoms of residue 554 are shown in stick style; carbon, nitrogen and oxygen atoms are colored white, blue and red, respectively. (d) Changes in molecular lipophilicity potential due to the G554V mutation, before and after energy minimization. The lipophilicity scores are mapped onto the molecular surface using a dark cyan (most hydrophilic) to dark goldenrod (most lipophilic) gradient. See **Fig. S2** for the full structural models, predicted local distance difference test and predicted template modelling scores. (e) Virus-like particles (VLPs) mL^−1^ in infected NIES-298 cultures from 0-24 hours post infection (hpi). Black, red and orange dots indicate the mean (*n* = 3) of gas-vacuolate (Gv) high N & high P (hN-hP), Gv low N & high P (lN-hP), and Gv low N & low P (lN-lP) groups, respectively. Error bars denote ± standard deviations.

The G554V mutation occurred not in the extracellular region of the protein exporter, but in its transmembrane domain. Therefore, we hypothesized that Ma-LMM01 would still be able to attach to but not infect resistant NIES-298 cells. Adsorption assays thus followed. For this experiment, we compared virion adsorption dynamics between the resistant gas-vacuolate NIES-298 variants in lN-hP and lN-lP media and susceptible gas-vacuolate cells in standard BG-11 (hN-hP) media as control, for 24 hours (the latent period of Ma-LMM01 lasts between 6-12 hours^33^). A clear drop in virus-like particles (VLPs) mL^−1^ was readily observed in susceptible and resistant groups throughout the latent period (0-6 hours post infection; hpi), indicative of attachment (**Fig. 3**e). No differences in VLPs mL^−1^ between groups were detected at any time point (1, 3, and 6 hpi) during the latent period (one-way ANOVA, *P*_adj_ ≥ 0.723 in all cases), suggesting similar attachment rates in susceptible and resistant NIES-298 subpopulations. As expected, VLPs increased at 24 hpi in the virus susceptible control and in the resistant groups the VLP concentration remained constant. At 24 hpi, we found that at least one group had a significant difference in VLPs mL^−1^ (log_10_) from one of the others (*F*_2,6_ = 58.53, *P*_adj_ = 4.64 × 10^−4^). We then used a Tukey’s test and confirmed that the difference was between susceptible and resistant groups (*P* ≤ 3.93 × 10^−4^ in both cases), and not between resistant subpopulations (*P* = 0.254). Collectively, these results indicated that Ma-LMM01 was able to attach to but not infect resistant NIES-298 cells.

### Gas vacuole infection experiment

We then infected susceptible gas-vacuolate and non-vacuolate NIES-298 with Ma-LMM01 in order to compare their infection dynamics in terms of growth rate (μ), Fv/Fm, phycocyanin (PC) content, and viral burst size. For this experiment, we used NIES-298 growing in standard (hN-hP) BG-11 media. Linear mixed-effects models (LMMs) detected no significant differences between uninfected gas-vacuolate and non-vacuolate subpopulations in μ and Fv/Fm (*P* ≥ 0.154 in both cases; **Fig. 4**a,b). In other words, uninfected subpopulations had a similar overall growth rate and photosynthetic efficiency. In contrast, μ and Fv/Fm in infected cultures differed significantly between gas-vacuolate and non-vacuolate groups (*P* ≤ 0.040 in both cases). In infected cultures, the mean growth rate of gas-vacuolate cells was lower (i.e., the mortality rate was higher) than that of non-vacuolate ones (**Fig 4**c). Accordingly, infected gas-vacuolate NIES-298 presented a lower mean Fv/Fm from 1-3 dpi compared to their non-vacuolate counterparts (**Fig. 4**e). This was consistent with the burst size of Ma-LMM01, which was significantly different between subpopulations at 1 dpi (Student’s *t*-test, *t*_4_ = 6.908, *P* = 0.002; **Fig. 4**d). Each gas-vacuolate cell produced 57.85 ± 5.59 viruses, while only 11.62 ± 6.06 viruses were produced per non-vacuolate cell. Finally, the PC/Chl *a* ratio was significantly higher in uninfected gas-vacuolate NIES-298 than in non-vacuolate groups (LMM, *P* = 0.011; **Fig. 4**f), whereas no difference was detected between subpopulations in infected cultures (*P* = 0.672; **Fig. 4**g).

**Fig. 4.**
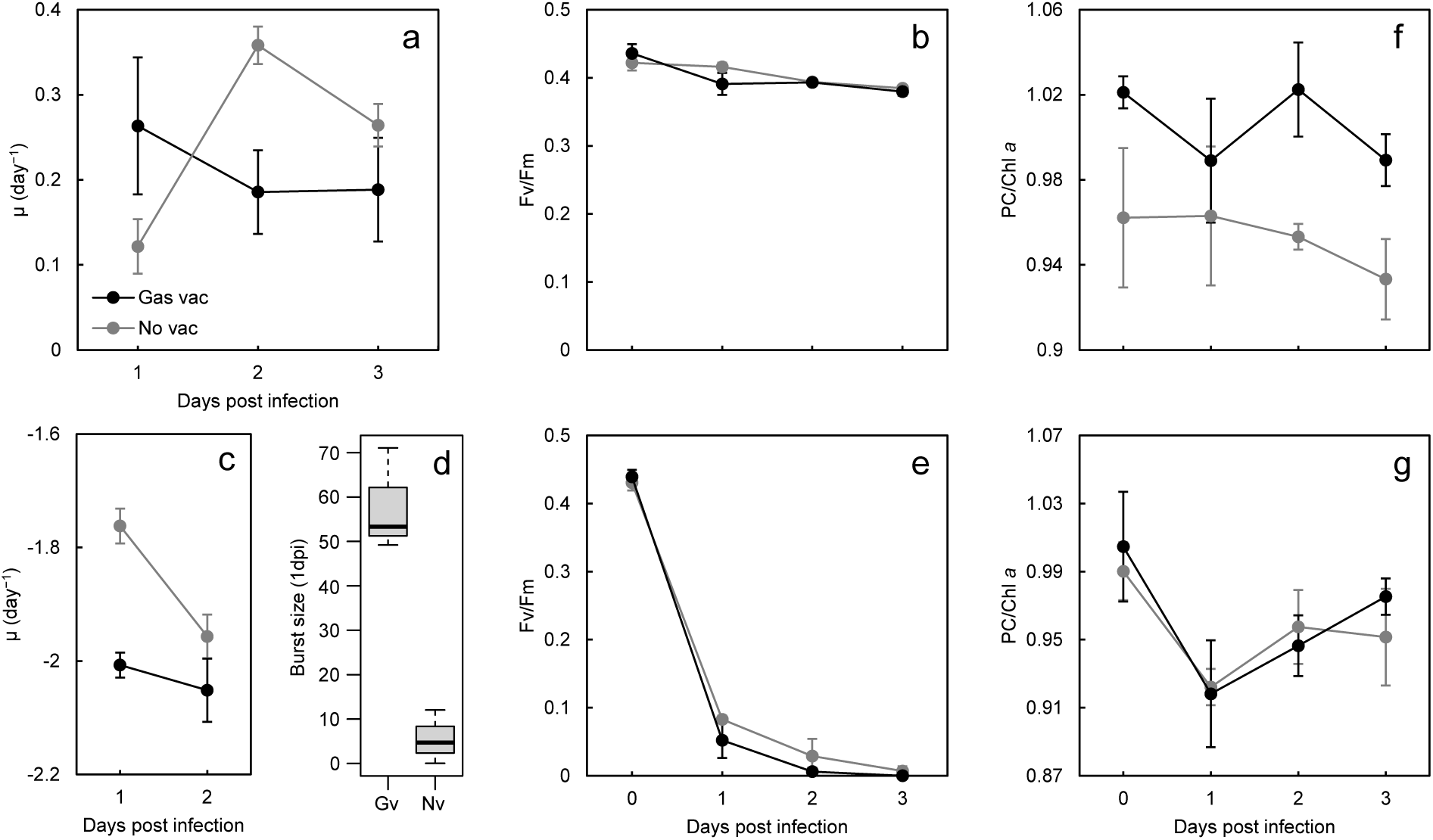
Gas vacuole infection experiment. (a) Growth rate (μ; day^−1^) and (b) Fv/Fm in uninfected *Microcystis aeruginosa* NIES-298 cultures over the course of the experiment. (c) μ up to 2 days post infection (dpi) in cultures infected with Ma-LMM01. (d) Burst size at 1 dpi (Gv, gas vacuole; Nv, no vacuole). (e) Fv/Fm in infected cultures. Phycocyanin (PC) to chlorophyll *a* (Chl *a*) ratio in (f) uninfected and (g) infected cultures. Black and gray dots indicate the mean (*n* = 3) of gas-vacuolate and non-vacuolate groups, respectively. Error bars denote ± standard deviations.

## Discussion

Eco-evolutionary processes involved in the acquisition of virus resistance have been relatively well studied in marine cyanobacteria^34–37^ and freshwater nitrogen fixers^28,29,38,39^. However, freshwater non-nitrogen-fixing cyanobacteria, such as the bloom-forming *M. aeruginosa*, have received comparatively little attention^30^, with research to date focusing only on temperature^27^. To the best of our knowledge, no studies have examined the evolution of virus resistance in non-nitrogen fixers within a nutrient and buoyancy experimental framework, despite the central roles of these factors in eutrophication and cyanoHABs in stratified lakes. In this work, we used the cyanophage Ma-LMM01 to experimentally investigate the acquisition of virus resistance in the toxic, non-nitrogen-fixer, freshwater, gas-vacuolate, bloom-forming cyanobacterium *M. aeruginosa* NIES-298 under different concentrations of nutrients (N and P).

After a year of adaptation to different nitrogen and phosphorus concentrations, resistance against Ma-LMM01 evolved specifically in gas-vacuolate NIES-298 cells under low N. This is in line with previous studies which have found that the availability of nitrogen shapes the evolution of phage resistance in nitrogen-fixing cyanobacteria^28,29^. In our study, three mutations were shared between resistant NIES-298 subpopulations (gas-vacuolate lN-hP and lN-lP) and absent in susceptible ones. Because these three mutations were present in both low N cultures and the lN-lP lineage was initially established from an lN-hP culture, they probably originated and were selected during the lN-hP adaptation period before being carried over and maintained in the lN-lP lineage. The first genetic variant occurred in an intergenic region far (199-259 bp) from any gene and was therefore deprioritized as a mutation conferring resistance. The second mutation took place in the amidophosphoribosyltransferase gene (*purF*) of NIES-298, where it changed a leucine to a valine. Both are hydrophobic amino acids, implying a small structural impact. PurF catalyzes the first committed step and is the rate-limiting enzyme of the *de novo* purine biosynthesis pathway^40,41^. Purine nitrogenous bases are N-rich compounds and, to the best of our knowledge, genetic changes in PurF or other purine biosynthesis enzymes have never been associated with virus resistance. Therefore, this SNP was likely related to adaptation to low nitrogen conditions and was not considered to confer cyanophage resistance. The final SNP unique to the resistant NIES-298 subpopulations occurred within a transmembrane protein. Mutations conferring cyanophage resistance have consistently been found to be predominantly located in membrane and cell surface-related proteins^28–30,34^. Hence, this genetic variant was further investigated.

The only membrane protein mutation unique to resistant (gas-vacuolate lN-hP and lN-lP) NIES-298 subpopulations occurred within the transmembrane domain of a peptidase domain-containing protein exporter. The SNP in this nitrogen-related transporter resulted in a glycine-to-valine substitution at position 554, a residue located right before the start of a predicted transmembrane α-helix. Glycine is known to facilitate local helix bending and affect the overall dynamics of transmembrane helices^42^. Accordingly, G554 in the wild type transporter was predicted to be inducing a bend in a helix. The G554V mutation substituted a small and flexible amino acid for a bulky and hydrophobic one, changing the original coil to a helix. The local hydrophobicity potential was also impacted; the molecular surface at this near-membrane position changed from hydrophilic to lipophilic, potentially promoting further membrane association. Collectively, the picture that emerged from both structural changes was one where the conformational flexibility of the transmembrane domain in the mutant transporter was negatively affected. Furthermore, the intracellular location of the peptidase domain suggests that this protein exporter degrades intracellular proteins and exports the resulting products to the extracellular environment. Because the G554V substitution occurred within the transmembrane domain rather than the peptidase domain, it may preserve intracellular protein degradation while impairing protein export. Under low nitrogen conditions, such a mutation could promote the intracellular retention of degradation products and potentially enhance nitrogen recycling within the cyanobacterial cell.

Since the mutation was located in the transmembrane domain of the protein rather than in its extracellular region, we hypothesized that Ma-LMM01 would still be able to attach to but not infect resistant NIES-298 cells. Adsorption assays indicated that this was indeed the case. In fact, a different membrane protein has previously been associated with Ma-LMM01 adsorption and resistance in NIES-298^30^. Together, these findings suggest that multiple host membrane proteins may be involved in Ma-LMM01 infection. Indeed, there are several bacteriophages that require more than one receptor for infection, such as Shigella phage Sf6 and the *Escherichia coli* phages T4 and T5^43^.

When we compared infection dynamics between susceptible gas-vacuolate and non-vacuolate NIES-298 subpopulations under the same conditions, we found that gas-vacuolate cells produced approximately five times more viruses than non-vacuolate cells. However, no overall differences in growth rate or Fv/Fm were detected between uninfected subpopulations, indicating that the overall fitness and photosynthetic efficiency of gas-vacuolate and non-vacuolate cells were similar. What caused then the drastic difference in burst size? We consider that a major contributing factor was the difference in PC content. PC is one of the main components of phycobilisomes, cyanobacterial light-harvesting complexes that can comprise up to half of the cellular soluble protein content^44–46^. PC was significantly higher in uninfected gas-vacuolate cells compared to their non-vacuolate counterparts. In contrast, no difference in PC content was found between infected subpopulations. This indicates that the additional PC and phycobilisomes of gas-vacuolate cells were degraded during infection. In accordance, Ma-LMM01 encodes the phycobilisome degradation protein NblA^32^, which is highly transcribed in the virocell^31,47.48^. Viral NblA proteins, including the NblA of Ma-LMM01^49^, have repeatedly been found to be more efficient at disassembling phycobilisomes than their cellular homologs^50–53^. The additional nitrogen that becomes available following virus-mediated phycobilisome and PC degradation provides further resources for viral synthesis during infection^31,32,49–52,54^, contributing to the larger burst size observed in gas-vacuolate cells.

Based on the experimental results presented here, we propose the following ecological model for the acquisition of virus resistance in stratified lakes during *Microcystis*-dominated blooms (**Fig. 5**). Gas-vacuolate *Microcystis* cells float and are therefore located in upper layers of the lake (relative to non-vacuolate cells), while non-vacuolate cells sink and are thus found deeper in the lake. Higher light intensities in upper layers lead to higher UV-induced DNA damage and mutation rates^55,56^. Lower nitrogen concentrations in upper layers^18^ lead to a stronger selective pressure to adapt to low nitrogen conditions, including adaptations in nitrogen-related transporters, such as protein exporters, which can be involved in virus infection (see also refs. 28,29). A larger viral burst size from gas-vacuolate *Microcystis* cells leads to more cyanophages, increased virus-host contact rates, more frequent infections and, consequently, a stronger selective pressure to evolve virus resistance in upper layers. As a result, compared to deeper layers, the upper layers of stratified lakes present a more conducive environment for the acquisition of virus resistance in *Microcystis*.

**Fig. 5.**
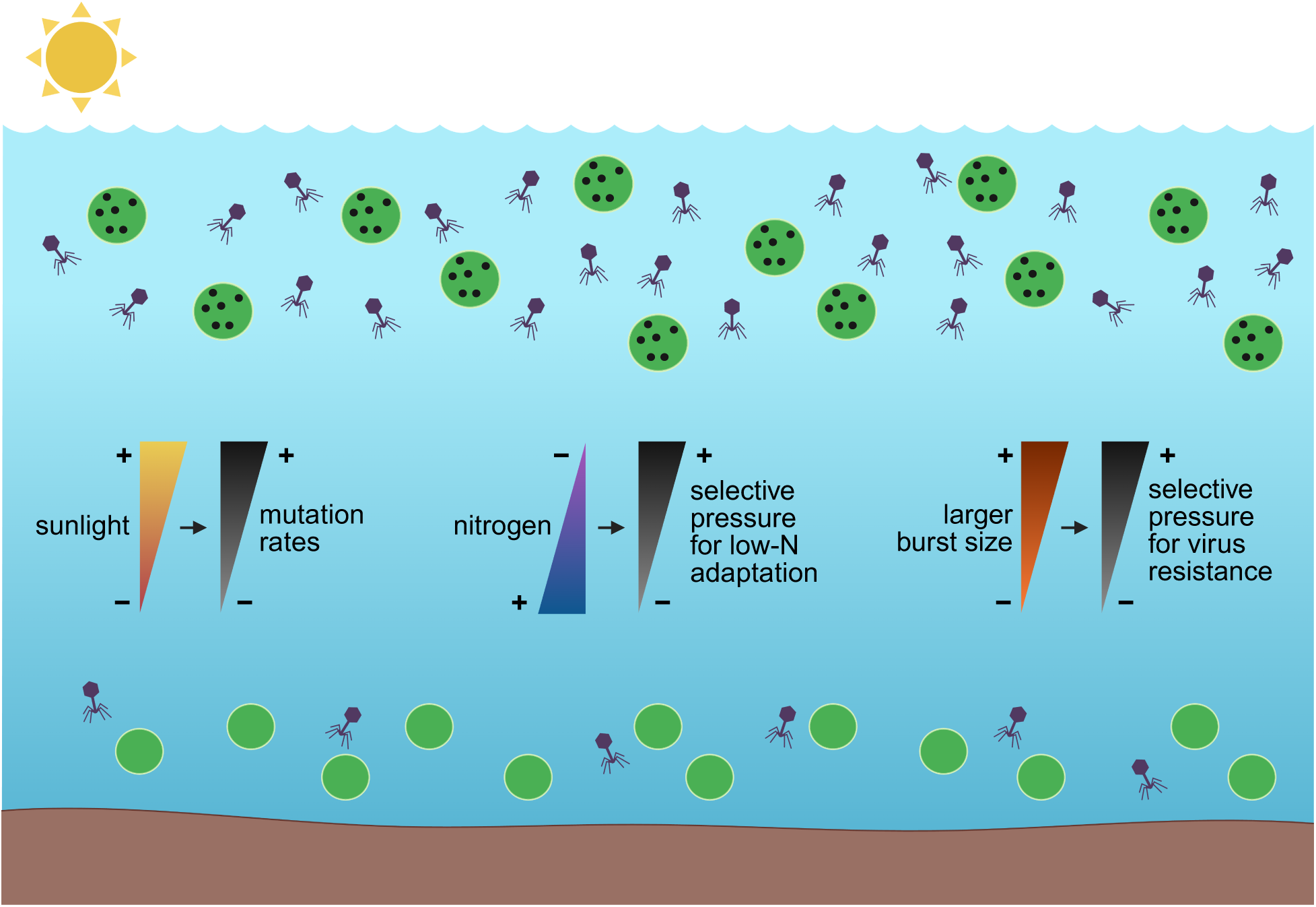
Experimentally derived ecological model for the acquisition of virus resistance in stratified lakes during *Microcystis*-dominated blooms. Gas-vacuolate *Microcystis* cells float and are therefore located in upper layers of the lake (relative to non-vacuolate cells), while non-vacuolate cells sink and are thus found deeper in the lake. Higher light intensities in upper layers lead to higher UV-induced DNA damage and mutation rates. Lower nitrogen concentrations in upper layers lead to a stronger selective pressure to adapt to low nitrogen conditions, including adaptations in nitrogen-related transporters, such as protein exporters, which can be involved in virus infection. A larger burst size from gas-vacuolate *Microcystis* cells leads to more cyanophages, increased virus-host contact rates, more frequent infections and, consequently, a stronger selective pressure to evolve virus resistance in upper layers. As a result, the upper layers of stratified lakes present a more favourable environment for the acquisition of virus resistance in *Microcystis* compared to deeper layers.

Overall, this work advances our understanding of how virus resistance can emerge and be structured within freshwater cyanobacterial populations. We show that under low nitrogen conditions, gas-vacuolate *M. aeruginosa* subpopulations acquire cyanophage resistance, resulting in pronounced differences in infection dynamics between buoyancy morphotypes. This suggests that nutrient availability can structure viral susceptibility within blooms by shaping the distribution of resistant and sensitive cells, potentially reducing the effectiveness of viral top-down control. Such nutrient-linked heterogeneity may therefore contribute to the persistence and internal restructuring of blooms rather than their collapse. More broadly, these findings highlight the importance of considering host physiology and vertical structure when interpreting virus-host dynamics in cyanobacterial blooms.

## Methods

### Cyanobacterial growth conditions and adaptation

*Microcystis aeruginosa* NIES-298, supplied axenic by the National Institute for Environmental Studies Microbial Culture Collection in Japan, was cultured in BG-11 media^57^ at 25 °C, 50 µmol m^−2^ s^−1^ and a light:dark cycle of 12:12 hours^58,59^. Four different concentrations of nitrogen and phosphorus were implemented in BG-11 in order to set up a nutrients matrix of two factors with two levels each. (1) 1500 mg L^−1^ NaNO_3_-N and 30 mg L^−1^ K_2_HPO_4_-P. This is the standard, full-strength BG-11, and it is referred to in the manuscript as high nitrogen and high phosphorus (abbreviated hN-hP). (2) 375 mg L^−1^ NaNO_3_-N, corresponding to 25% of the standard BG-11 NaNO_3_-N concentration, and 30 mg L^−1^ K_2_HPO_4_-P. This modified version of BG-11 is referred here as low nitrogen and high phosphorus (lN-hP). (3) 1500 mg L^−1^ NaNO_3_-N and 7.5 mg L^−1^ K_2_HPO_4_-P, which corresponds to 25% of the standard BG-11 K_2_HPO_4_-P concentration. This BG-11 version is referred to as high nitrogen and low phosphorus (hN-lP). And (4) 375 mg L^−1^ NaNO_3_-N and 7.5 mg L^−1^ K_2_HPO_4_-P, which is referred to as low nitrogen and low phosphorus BG-11 (lN-lP). The remaining nutrients were left unchanged in all BG-11 versions. NIES-298 was first propagated concurrently from hN-hP BG-11 to lN-hP and hN-lP concentrations. Subsequently, an lN-hP-adapted NIES-298 culture was used as the inoculum to adapt the cyanobacteria to lN-lP BG-11. The cultures were grown in these conditions throughout a year, subculturing them every two to three weeks into fresh media^60^.

### Cyanobacterial and viral measurements

Cyanobacterial cell density in all experiments was quantified in a FACSAria Fusion Flow Cytometer (BD) equipped with a 488 nm laser by analyzing live samples using red fluorescence vs side scatter. FC data analysis and generation of plots were conducted in FlowJo 10.10.0. The maximum quantum yield of photosystem II (Fv/Fm) was monitored as a measure of the photosynthetic efficiency of the cyanobacterial cells. For this, 20 µL of the cultures were diluted in 1980 µL of BG-11, acclimated in the dark for 10 minutes, and measured in a WATER-PAM chlorophyll fluorometer (Heinz Walz GmbH). Chlorophyll *a* (Chl *a*) and phycocyanin (PC) were measured in a Synergy LX (BioTek) microplate reader using whole-cell absorbance at 680 nm and 630 nm, respectively^50,53^. PC and Chl *a* absorbances were then divided to normalize PC against Chl *a* and obtain a PC/Chl *a* ratio^53^.

Infectious virus titre was quantified using the most probable number (MPN) protocol described previously^58^, which is based on the original method developed in ref. 61. Briefly, NIES-298 in exponential growth phase (∼5 × 10⁵ cells mL^−1^) was infected with 10-fold serial dilutions of 0.45 μm syringe-filtered Ma-LMM01 lysate in a 96-well plate. The plates were visually inspected for lysis and scored five dpi. The MPN was then calculated using the method in ref. 62 implemented in the MPN package in R. VLPs were measured via flow virometry (FV) using the protocol developed in refs. 63-65. In brief, 10 μL of 25% glutaraldehyde was added to 500 μL of sample in cryotubes before incubating them at 4 °C for 15 minutes. The cryotubes were then snap-frozen in liquid nitrogen and stored at −80 °C until analysis. Samples were thawed at room temperature and diluted in TE buffer (pH 8) by adding 100 µL of sample to 900 µL of TE. After staining with 10 µL of SYBR Green I working solution (5 µL of dye stock in 995 µL of 0.02 µm syringe-filtered Milli-Q water), samples were heated at 80 °C for 10 minutes in the dark and subsequently cooled down at room temperature for 5 minutes. The samples were then analyzed in a FACSAria Fusion Flow Cytometer using green fluorescence vs side scatter.

### Infection experiments

For the nutrients infection experiment (**Fig. 1**), exponentially growing NIES-298 cultures adapted to the four different BG-11 nutrient concentrations (hN-hP, lN-hP, hN-lP, and lN-lP) were infected with Ma-LMM01^33^ as follows. 150 mL master cultures (one for each nutrient concentration) were split into six replicates^58,59^ and three cultures were infected with 0.45 μm syringe-filtered Ma-LMM01 lysate. The other three cultures were mock-infected with autoclaved Ma-LMM01 lysate and served as uninfected controls. Cyanobacterial cell density and Fv/Fm were monitored daily until 3 dpi as described above.

For the confirmatory (resistant vs susceptible cells; **Fig. S1**) infection experiment, gas-vacuolate and non-vacuolate hN-hP, lN-hP, hN-lP, and lN-lP NIES-298 subpopulations in exponential phase were infected with Ma-LMM01. 150 mL master cultures were split into six replicates, and three cultures were infected with 0.45 μm syringe-filtered Ma-LMM01. Autoclaved Ma-LMM01 lysate was added to the other three cultures as uninfected controls. Cyanobacterial cell density was measured at 0 dpi and 7 dpi.

For the gas vacuole infection experiment (**Fig. 4**), NIES-298 cultures in exponential phase containing gas-vacuolate or non-vacuolate cells in standard (hN-hP) BG-11 were infected as follows. 150 mL master cultures were split into six replicates, and three cultures were infected with 0.45 μm syringe-filtered Ma-LMM01 lysate. The control uninfected cultures were spiked with autoclaved Ma-LMM01 lysate. Cyanobacterial cell density, Fv/Fm, infectious viruses and PC/Chl *a* ratio were monitored every day until 3 dpi as described above. All infection experiments were conducted at an average multiplicity of infection (MOI) of 3.49 ± 0.97 viruses cell^−1^.

### Cell sorting

Cell sorting of gas-vacuolate and non-vacuolate NIES-298 subpopulations growing in hN-hP, lN-hP, hN-lP, and lN-lP BG-11 was conducted using a FACSAria Fusion Flow Cytometer (BD). After preparing the instrument for aseptic sorting with standard protocols as per the manufacturer, the two morphotypes were sorted based on their side scatter, forward scatter, and red fluorescence properties. 1 mL of sorted exponentially growing NIES-298 cultures was received in 12.5 mL of the corresponding BG-11 media contained in sterile 15 mL Falcon tubes. For example, NIES-298 growing in hN-hP was sorted into hN-hP BG-11, lN-hP cells into lN-hP media, and so on. After sorting, the 13.5 mL were immediately transferred to sterile 50 mL Cellstar flasks (Greiner Bio-One) and cultured under the conditions described earlier. The sorted cultures were then carefully examined under a 40X objective lens using a Revolve (ECHO) microscope in brightfield mode.

### Adsorption assays

For virus adsorption assays, resistant (gas-vacuolate lN-hP and lN-lP) and susceptible (gas-vacuolate hN-hP) exponentially growing NIES-298 cultures were infected with Ma-LMM01 as follows: 75 mL master cultures were divided into three replicates and infected with 0.45 μm syringe-filtered Ma-LMM01 lysate at a VLP-to-cell ratio of 0.08 ± 0.01. VLPs were quantified by FV at 1, 3, 6, and 24 hpi as described previously.

### DNA extraction and sequencing

In order to concentrate cyanobacterial cells for DNA extraction, 50 mL of gas-vacuolate and non-vacuolate NIES-298 cultures (hN-hP, lN-hP, hN-lP, and lN-lP) in exponential phase were filtered through 0.45 μm Millex-HP (Millipore) syringe filters. Once the cells were concentrated on the filters, 25 mL of molecular biology grade water (Corning) was passaged through the filters to rinse any residual BG-11 compounds that could interfere with downstream sample processing. DNA extraction from filters and purification were conducted with a MasterPure Complete DNA and RNA Purification Kit (Biosearch Technologies) using the protocol developed in refs. 66,67. RNA was removed by enzymatic digestion with RNase A supplied in the kit. Library preparation and paired-end sequencing were carried out at the Waterloo Genomics Surveillance Centre with an Illumina DNA Prep kit using standard methods as per the manufacturer, a read length of 300 bp, and a NextSeq 2000 (Illumina) sequencer.

### Genomics pipeline

Quality assessment and filtering of the raw reads were conducted using FastQC 0.12.1^68^ and fastp 1.0.1^69^, respectively. Read mapping and variant calling were performed using *breseq* 0.38.2^70,71^ with Bowtie 2 2.5.2^72^ and R 4.5.0^73^. The complete reference genome of NIES-298 (RefSeq accession number: NZ_CP184724.1) and its plasmids (NZ_PV068373.1 and NZ_PV076616.1) sequenced in ref. 74 were used for alignment. Detected mutations were manually inspected^28,29^ using the native HTML output files and BAM files generated by *breseq*, with the latter examined in Tablet 1.21.02.08^75^.

### Structural bioinformatics workflow

Structural modelling was conducted using AlphaFold 3 (AF3)^76^. The overall fold of AF3 models with predicted template modelling (pTM) scores > 0.5 is usually considered similar to that of the true structure^77,78^. Since the wildtype and mutant protein exporter models had pTM scores = 0.73, they were used for downstream analyses. Additionally, the predicted local distance difference test scores of the AF3 models were visually inspected in the native environment of the AlphaFold server and can be found in **Fig. S2**. Energy minimization was performed in ChimeraX 1.12^79^ using Antechamber^80^ and OpenMM 8^81^. The molecular lipophilicity potential was calculated in ChimeraX as well. Transmembrane regions and topology were predicted using DeepTMHMM 1.0^82^. Molecular visualization was carried out in PyMol 3.1.8^83^, and Missense3D^84^ was used to corroborate the impacts of amino acid substitutions. Domains were identified using InterProScan^85^ and domain architecture schematics were elaborated with IBS 2.0^86^.

### Statistical analyses

Statistical analyses were conducted using R 4.6.0^73^. In order to investigate whether there was a significant interaction between N and P on cyanobacterial cell density and photosynthetic efficiency (Fv/Fm) in NIES-298 cultures infected with Ma-LMM01 during the last day of the nutrients infection experiment, two-way ANOVAs were employed. Prior to analyses, the normality of residuals and the homogeneity of variances were tested using Shapiro-Wilk and Levene’s (car package) tests, respectively. Fv/Fm residuals were normally distributed (*P* = 0.206) and the variances were homogeneous (*P* = 0.929). Cell density residuals, however, departed from normality. Thus, a log_10_ transformation was applied to cell density data. After transformation, cell density residuals followed normal distributions (*P* = 0.925) and the variances were homogeneous (*P* = 0.608). In these analyses, N and P were used as two-level (low & high) factors. Fv/Fm and log_10_-transformed cell density values were used as dependent variables. Further differences were investigated using Tukey’s *a posteriori* tests.

To test for differences in the percentage change in cell density of NIES-298 subpopulations in lN-hP and lN-lP cultures from 0 dpi to 3 dpi during the nutrients infection experiment, we used paired *t*-tests. The normality of differences was investigated using Shapiro-Wilk tests before analyses. The differences followed normal distributions (*P* ≥ 0.197 in both cases). Subpopulation (weaker side scatter vs stronger side scatter) was used as the grouping variable and percentage change in cell density as the dependent variable.

In order to investigate differences in VLPs mL^−1^ between gas-vacuolate hN-hP, lN-hP, and lN-lP NIES-298 cultures infected with Ma-LMM01 at 1, 3, 6, and 24 hpi, we performed separate one-way ANOVAs for each time point and the resulting *P*-values were adjusted using the Benjamini-Hochberg false discovery rate correction. The residuals at 1, 3, and 6 hpi were normally distributed (Shapiro-Wilk test, *P* ≥ 0.470) and variances were homogeneous (Levene’s test, *P* ≥ 0.399). However, VLPs residuals at 24 hpi departed from normality. A log_10_ transformation was therefore applied to VLPs data at 24 hpi. After transformation, the VLPs residuals at 24 hpi followed a normal distribution (*P* = 0.398) and the variances were homogeneous (*P* = 0.404). Tukey’s tests were used to further investigate significant differences.

Linear mixed-effects models were employed to investigate overall differences in growth rate (μ), Fv/Fm, and PC/Chl *a* ratio between gas-vacuolate and non-vacuolate NIES-298 subpopulations throughout the infection experiment (lme4 and lmerTest packages). The normality and homoscedasticity of residuals were assessed visually using QQ and residual plots and statistically using Shapiro-Wilk and Breusch-Pagan (lmtest package) tests. No deviations from normality or homoscedasticity were visually observed or statistically detected (Shapiro-Wilk, *P* ≥ 0.086; Breusch-Pagan, *P* ≥ 0.245 in all cases). Group (gas-vacuolate, non-vacuolate) and dpi (0-3) were treated as categorical variables, and Fv/Fm, μ (day^−1^) and PC/Chl *a* ratio were used as response variables. Because most cells had lysed by 2 dpi, only μ up to 2 dpi was calculated for the infected groups. Replicate identity was included as a random intercept to account for repeated measurements over time. Finally, a Student’s *t*-test was used to investigate whether there was a significant difference in the burst size of Ma-LMM01 between gas-vacuolate and non-vacuolate NIES-298 subpopulations at 1 dpi. The data followed normal distributions (Shapiro-Wilk test, *P* ≥ 0.339 in both cases) and the variances were homogeneous (Levene’s test, *P* = 0.601). Significance was defined at *P* < 0.05.

## Acknowledgements

This work was funded by a Grant-in-Aid of Research (project: Genomic insights into virus resistance in toxic cyanobacteria) by the Phycological Society of America (PSA), an international Ontario Graduate Scholarship (OGS) and a Secretaría de Ciencia, Humanidades, Tecnología e Innovación (SECIHTI) Becas de Posgrado para Maestrías y Doctorados en Ciencias y Humanidades en el Extranjero scholarship awarded to I.M.P.; and Natural Sciences and Engineering Research Council of Canada grants (RGPIN-2022-03350 and DGECR-2022-00329) and a 2021 Phycological Society of America Norma J. Lang Early Career Research Fellowship awarded to J.I.N.

We would also like to thank Prof. Takashi Yoshida for providing the cyanophage Ma-LMM01 as a kind gift; Akash Jairaj at the UW Molecular Biology Core Facility for valuable technical advice on cell sorting and flow cytometry; Dr. Nicole Poulton at the Bigelow Laboratory for Ocean Sciences for an excellent course on aquatic flow cytometry; Prof. Kirsten M. Müller for kindly letting us use her microscope; Heather Roshon at the Canadian Phycological Culture Centre for her help with media preparation; Hadi Dhiyebi and Hoang Dang at the Waterloo Genomics Surveillance Centre for their help with library prep and sequencing; and the Digital Research Alliance of Canada for their computational resources and support during the bioinformatics work.

## Author contributions

I.M.P. and J.I.N. conceived the ideas and acquired funding. I.M.P performed the experiments, sorted the cells, conducted the bioinformatics pipeline, analyzed the data, conceptualized the ecological model, and wrote the manuscript. J.I.N supervised the work. Both authors revised and edited the manuscript.

