## Supplementary Material for "Gas-vacuolate *Microcystis* evolves cyanophage resistance under low nitrogen conditions"

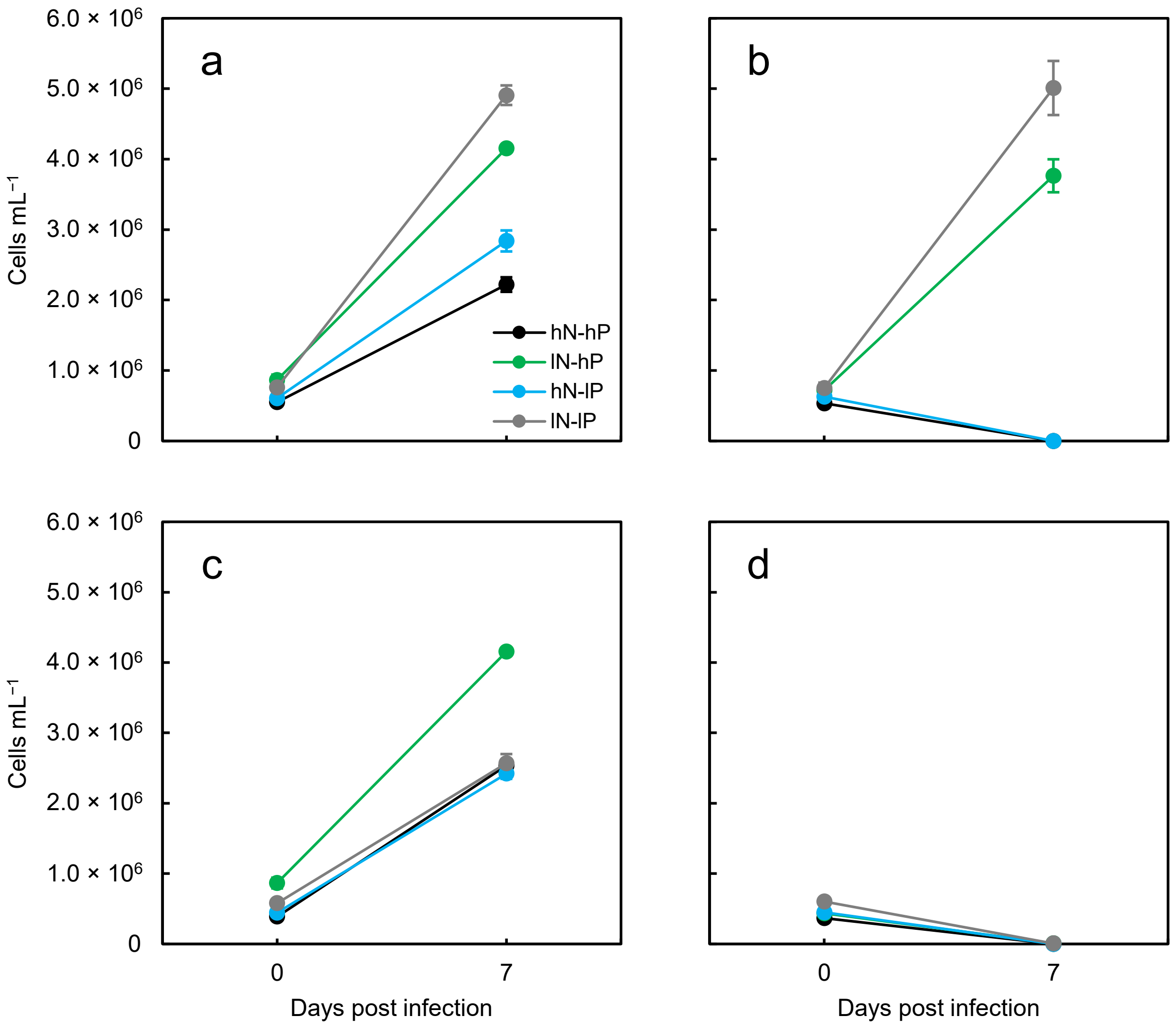


**Fig. S1**. Confirmatory infection experiment. (a) Cells mL$^{-1}$ in uninfected gas-vacuolate *Microcystis aeruginosa* NIES-298 cultures over the course of the experiment, and (b) cells mL$^{-1}$ in gas-vacuolate cultures infected with Ma-LMM01. (c) Cells mL$^{-1}$ in uninfected non-vacuolate cultures, (d) cells mL$^{-1}$ in infected non-vacuolate cultures. Black, green, light blue, and gray dots indicate the mean ($n$ = 3) of high N & high P (hN-hP), low N & high P (lN-hP), high N & low P (hN-lP), and low N & low P (lN-lP) groups, respectively. Error bars denote ± standard deviations.

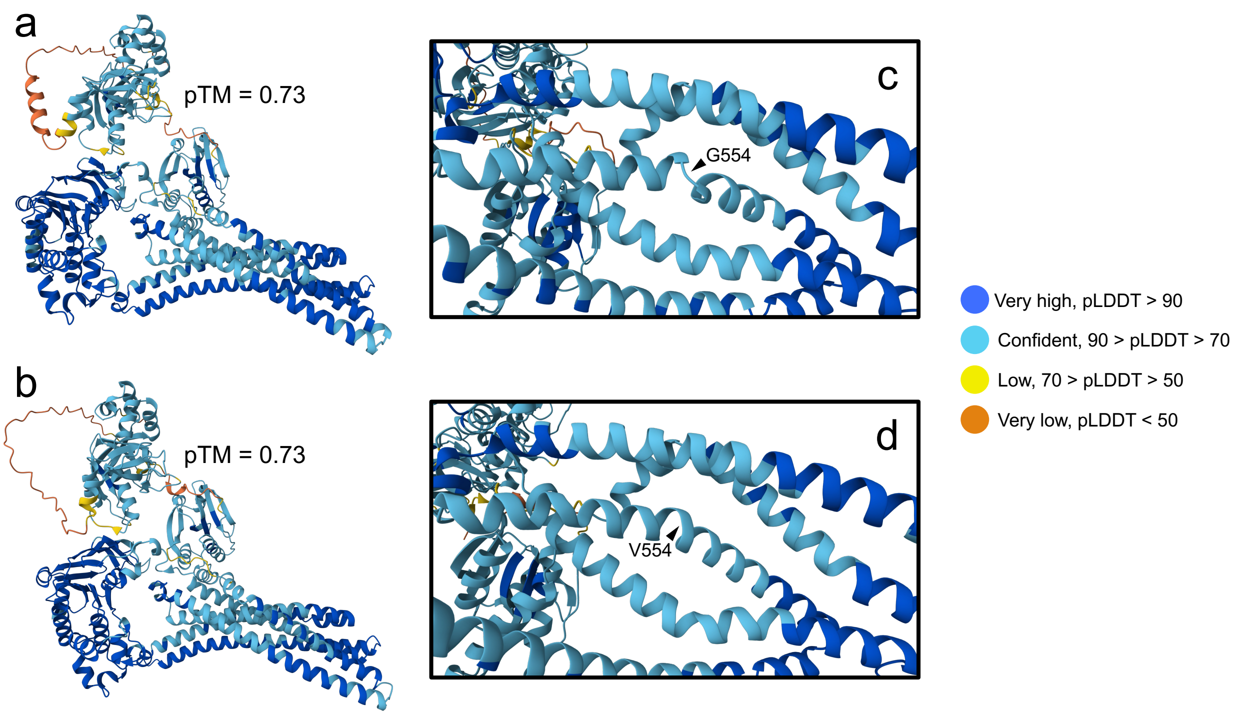


**Fig. S2**. Predicted local distance difference test (pLDDT) and predicted template modelling (pTM) scores of the AlphaFold 3 structural models. (a) Wild type protein exporter of *Microcystis aeruginosa* NIES-298. (b) Mutant protein exporter. (c,d) Close-up views into the mutant residue region for wild type and mutant transporters, respectively. The mutant residue (glycine in wild type and valine in mutant) is labeled and highlighted with an arrow.

**Table S1**. Mutations in gas-vacuolate and non-vacuolate *Microcystis aeruginosa* NIES-298 subpopulations. Mutations shared between resistant NIES-298 subpopulations and absent in susceptible ones are highlighted in bold. Gas vacuole-related mutations are italicized.

| Population | Location | Position | Mutation[a] | Description | Gene | Annotation | Frequency[b] |
|---|---|---|---|---|---|---|---|
| hN-hP, Gv | Chr | 568,519 | C→A | noncoding (1282/1489 nt) | ACNQK5_RS02750 ← | 16S ribosomal RNA | 100, 10 |
| hN-hP, Gv | Chr | 568,530 | G→T | noncoding (1271/1489 nt) | ACNQK5_RS02750 ← | 16S ribosomal RNA | 100, 11 |
| hN-hP, Gv | Chr | 568,541 | C→A | noncoding (1260/1489 nt) | ACNQK5_RS02750 ← | 16S ribosomal RNA | 100, 11 |
| hN-hP, Gv | Chr | 568,560 | C→T | noncoding (1241/1489 nt) | ACNQK5_RS02750 ← | 16S ribosomal RNA | 100, 9 |
| hN-hP, Gv | Chr | 568,568 | Δ1 bp | noncoding (1233/1489 nt) | ACNQK5_RS02750 ← | 16S ribosomal RNA | 100, 9 |
| hN-hP, Gv | Chr | 568,577 | C→G | noncoding (1224/1489 nt) | ACNQK5_RS02750 ← | 16S ribosomal RNA | 100, 9 |
| hN-hP, Gv | Chr | 2,023,374 | (AAAGGCA)15→19 | coding (118/243 nt) | ACNQK5_RS09745 → | hypothetical protein | 86.6, 282 |
| hN-hP, Gv | Chr | 2,367,866 | Δ144 bp | coding (3-146/213 nt) | ACNQK5_RS11275 ← | hypothetical protein | 100, 322 |
| *hN-hP, Gv* | *Chr* | *2,461,108* | *A→G* | *intergenic (-215/+53)* | *gvpC ← / ← gvpA* | *gas vesicle protein GvpC/gas vesicle structural protein GvpA* | *89.8, 215* |
| *hN-hP, Gv* | *Chr* | *2,461,116* | *2 bp→CA* | *intergenic (-223/+44)* | *gvpC ← / ← gvpA* | *gas vesicle protein GvpC/gas vesicle structural protein GvpA* | *89.3, 206* |
| *hN-hP, Gv* | *Chr* | *2,461,128* | *A→C* | *intergenic (-235/+33)* | *gvpC ← / ← gvpA* | *gas vesicle protein GvpC/gas vesicle structural protein GvpA* | *88.8, 188* |
| *hN-hP, Gv* | *Chr* | *2,461,132* | *G→C* | *intergenic (-239/+29)* | *gvpC ← / ← gvpA* | *gas vesicle protein GvpC/gas vesicle structural protein GvpA* | *87.7, 179* |
| *hN-hP, Gv* | *Chr* | *2,461,140* | *A→G* | *intergenic (-247/+21)* | *gvpC ← / ← gvpA* | *gas vesicle protein GvpC/gas vesicle structural protein GvpA* | *87.9, 175* |
| *hN-hP, Gv* | *Chr* | *2,461,144* | *T→G* | *intergenic (-251/+17)* | *gvpC ← / ← gvpA* | *gas vesicle protein GvpC/gas vesicle structural protein GvpA* | *87.9, 173* |
| *hN-hP, Gv* | *Chr* | *2,461,155* | *2 bp→GG* | *intergenic (-262/+5)* | *gvpC ← / ← gvpA* | *gas vesicle protein GvpC/gas vesicle structural protein GvpA* | *84.2, 159* |
| *hN-hP, Gv* | *Chr* | *2,461,158* | *2 bp→TT* | *intergenic (-265/+2)* | *gvpC ← / ← gvpA* | *gas vesicle protein GvpC/gas vesicle structural protein GvpA* | *84.4, 162* |
| *hN-hP, Gv* | *Chr* | *2,461,176* | *T→C* | *A67A (GCA→GCG)* | *gvpA ←* | *gas vesicle structural protein GvpA* | *81.7, 144* |
| *hN-hP, Gv* | *Chr* | *2,461,633* | *Δ1 bp* | *intergenic (-257/+73)* | *gvpA ← / ← gvpA* | *gas vesicle structural protein GvpA/gas vesicle structural protein GvpA* | *80.4, 102* |
| hN-hP, Gv | Chr | 3,397,270 | C→T | noncoding (1063/1489 nt) | ACNQK5_RS16330 → | 16S ribosomal RNA | 83.3, 12 |
| hN-hP, Gv | Chr | 3,397,273 | T→C | noncoding (1066/1489 nt) | ACNQK5_RS16330 → | 16S ribosomal RNA | 100, 10 |
| hN-hP, Gv | Chr | 3,397,284 | +C | noncoding (1077/1489 nt) | ACNQK5_RS16330 → | 16S ribosomal RNA | 80.0, 10 |
| hN-hP, Gv | Chr | 3,397,355 | G→T | noncoding (1148/1489 nt) | ACNQK5_RS16330 → | 16S ribosomal RNA | 100, 12 |
| hN-hP, Gv | Chr | 3,397,360 | C→G | noncoding (1153/1489 nt) | ACNQK5_RS16330 → | 16S ribosomal RNA | 100, 12 |
| hN-hP, Gv | Chr | 3,397,369 | T→G | noncoding (1162/1489 nt) | ACNQK5_RS16330 → | 16S ribosomal RNA | 100, 12 |
| hN-hP, Gv | Chr | 3,397,372 | T→A | noncoding (1165/1489 nt) | ACNQK5_RS16330 → | 16S ribosomal RNA | 83.3, 12 |
| hN-hP, Gv | Chr | 3,397,377 | G→T | noncoding (1170/1489 nt) | ACNQK5_RS16330 → | 16S ribosomal RNA | 100, 12 |

| | | | | | | | |
|---|---|---|---|---|---|---|---|
| hN-hP, Gv | Chr | 3,397,404 | A→G | noncoding (1197/1489 nt) | ACNQK5_RS16330 → | 16S ribosomal RNA | 100, 12 |
| hN-hP, Gv | Chr | 3,397,417 | T→C | noncoding (1210/1489 nt) | ACNQK5_RS16330 → | 16S ribosomal RNA | 83.3, 12 |
| hN-hP, Gv | Chr | 3,397,424 | A→G | noncoding (1217/1489 nt) | ACNQK5_RS16330 → | 16S ribosomal RNA | 100, 12 |
| hN-hP, Gv | Chr | 3,397,431 | G→C | noncoding (1224/1489 nt) | ACNQK5_RS16330 → | 16S ribosomal RNA | 83.3, 12 |
| hN-hP, Gv | Chr | 3,397,440 | Δ1 bp | noncoding (1233/1489 nt) | ACNQK5_RS16330 → | 16S ribosomal RNA | 100, 12 |
| hN-hP, Gv | Chr | 3,397,448 | 2 bp→AT | noncoding (1241-1242/1489 nt) | ACNQK5_RS16330 → | 16S ribosomal RNA | 83.3, 12 |
| hN-hP, Gv | Chr | 3,397,458 | A→G | noncoding (1251/1489 nt) | ACNQK5_RS16330 → | 16S ribosomal RNA | 100, 13 |
| hN-hP, Gv | Chr | 3,397,467 | G→T | noncoding (1260/1489 nt) | ACNQK5_RS16330 → | 16S ribosomal RNA | 100, 14 |
| hN-hP, Gv | Chr | 3,397,478 | C→A | noncoding (1271/1489 nt) | ACNQK5_RS16330 → | 16S ribosomal RNA | 100, 15 |
| hN-hP, Gv | Chr | 3,397,489 | 2 bp→TC | noncoding (1282-1283/1489 nt) | ACNQK5_RS16330 → | 16S ribosomal RNA | 100, 15 |
| hN-hP, Gv | Chr | 3,466,568 | C→A | A617A (GCG→GCT) | ACNQK5_RS16575 ← | beta strand repeat-containing protein | 100, 47 |
| hN-hP, Gv | Chr | 3,466,571 | G→A | D616D (GAC→GAT) | ACNQK5_RS16575 ← | beta strand repeat-containing protein | 100, 47 |
| hN-hP, Gv | P2 | 40,585 | A→G | pseudogene (241/732 nt) | PLSM73337038_RS00215 → | IS5-like element ISMae4 family transposase | 91.6, 214 |
| hN-hP, Gv | P2 | 40,793 | T→C | pseudogene (449/732 nt) | PLSM73337038_RS00215 → | IS5-like element ISMae4 family transposase | 90.2, 909 |
| hN-hP, Nv | Chr | 339,532 | (ACCTAT)1→2 | pseudogene (403/423 nt) | ACNQK5_RS01640 ← | transposase family protein | 91.2, 338 |
| hN-hP, Nv | Chr | 2,023,374 | (AAAGGCA)15→19 | coding (118/243 nt) | ACNQK5_RS09745 → | hypothetical protein | 86.0, 344 |
| hN-hP, Nv | Chr | 2,367,866 | Δ144 bp | coding (3-146/213 nt) | ACNQK5_RS11275 ← | hypothetical protein | 100, 381 |
| hN-hP, Nv | Chr | 3,397,355 | G→T | noncoding (1148/1489 nt) | ACNQK5_RS16330 → | 16S ribosomal RNA | 100, 7 |
| hN-hP, Nv | Chr | 3,397,377 | G→T | noncoding (1170/1489 nt) | ACNQK5_RS16330 → | 16S ribosomal RNA | 100, 7 |
| hN-hP, Nv | Chr | 3,397,440 | Δ1 bp | noncoding (1233/1489 nt) | ACNQK5_RS16330 → | 16S ribosomal RNA | 87.5, 8 |
| hN-hP, Nv | Chr | 3,397,458 | A→G | noncoding (1251/1489 nt) | ACNQK5_RS16330 → | 16S ribosomal RNA | 88.9, 9 |
| hN-hP, Nv | Chr | 3,397,478 | C→A | noncoding (1271/1489 nt) | ACNQK5_RS16330 → | 16S ribosomal RNA | 88.9, 9 |
| hN-hP, Nv | Chr | 3,466,568 | C→A | A617A (GCG→GCT) | ACNQK5_RS16575 ← | beta strand repeat-containing protein | 97.6, 41 |
| hN-hP, Nv | Chr | 3,466,571 | G→A | D616D (GAC→GAT) | ACNQK5_RS16575 ← | beta strand repeat-containing protein | 97.4, 39 |
| hN-hP, Nv | P2 | 40,585 | A→G | pseudogene (241/732 nt) | PLSM73337038_RS00215 → | IS5-like element ISMae4 family transposase | 94.7, 281 |
| hN-hP, Nv | P2 | 40,793 | T→C | pseudogene (449/732 nt) | PLSM73337038_RS00215 → | IS5-like element ISMae4 family transposase | 93.7, 1052 |
| lN-hP, Gv | Chr | 2,023,374 | (AAAGGCA)15→19 | coding (118/243 nt) | ACNQK5_RS09745 → | hypothetical protein | 86.7, 391 |
| lN-hP, Gv | Chr | 2,367,866 | Δ144 bp | coding (3-146/213 nt) | ACNQK5_RS11275 ← | hypothetical protein | 100, 369 |
| *lN-hP, Gv* | *Chr* | *2,462,329* | *G→T* | *L55I (CTC→ATC)* | *gvpA ←* | *gas vesicle structural protein GvpA* | *100, 433* |
| *lN-hP, Gv* | *Chr* | *2,462,342* | *A→C* | *S50S (TCT→TCG)* | *gvpA ←* | *gas vesicle structural protein GvpA* | *100, 440* |

| | | | | | | | |
|---|---|---|---|---|---|---|---|
| **lN-hP, Gv** | **Chr** | **3,210,317** | **(C)12→11** | **intergenic (-199/-259)** | **ACNQK5_RS15405 ← / → ACNQK5_RS15410** | **PEP-CTERM sorting domain-containing protein/Uma2 family endonuclease** | **96.0, 178** |
| lN-hP, Gv | Chr | 3,466,568 | C→A | A617A (GCG→GCT) | ACNQK5_RS16575 ← | beta strand repeat-containing protein | 91.8, 49 |
| lN-hP, Gv | Chr | 3,466,571 | G→A | D616D (GAC→GAT) | ACNQK5_RS16575 ← | beta strand repeat-containing protein | 93.5, 46 |
| **lN-hP, Gv** | **Chr** | **3,802,561** | **G→T** | **G554V (GGT→GTT)** | **ACNQK5_RS18290 →** | **peptidase domain-containing ABC transporter** | **99.5, 390** |
| **lN-hP, Gv** | **Chr** | **4,213,818** | **G→C** | **L71V (CTT→GTT)** | **purF ←** | **amidophosphoribosyltransferase** | **99.8, 458** |
| lN-hP, Gv | P2 | 40,793 | T→C | pseudogene (449/732 nt) | PLSM73337038_RS00215 → | IS5-like element ISMae4 family transposase | 81.8, 1986 |
| lN-hP, Nv | Chr | 568,330 | A→G | noncoding (1471/1489 nt) | ACNQK5_RS02750 ← | 16S ribosomal RNA | 100, 10 |
| lN-hP, Nv | Chr | 568,340 | G→A | noncoding (1461/1489 nt) | ACNQK5_RS02750 ← | 16S ribosomal RNA | 100, 10 |
| lN-hP, Nv | Chr | 568,374 | 3 bp→TCG | noncoding (1425-1427/1489 nt) | ACNQK5_RS02750 ← | 16S ribosomal RNA | 100, 10 |
| lN-hP, Nv | Chr | 568,380 | 3 bp→ACA | noncoding (1419-1421/1489 nt) | ACNQK5_RS02750 ← | 16S ribosomal RNA | 100, 10 |
| lN-hP, Nv | Chr | 568,385 | T→G | noncoding (1416/1489 nt) | ACNQK5_RS02750 ← | 16S ribosomal RNA | 100, 10 |
| lN-hP, Nv | Chr | 568,387 | Δ1 bp | noncoding (1414/1489 nt) | ACNQK5_RS02750 ← | 16S ribosomal RNA | 90, 10 |
| lN-hP, Nv | Chr | 568,392 | 1 bp→AG | noncoding (1409/1489 nt) | ACNQK5_RS02750 ← | 16S ribosomal RNA | 90, 10 |
| lN-hP, Nv | Chr | 568,394 | C→G | noncoding (1407/1489 nt) | ACNQK5_RS02750 ← | 16S ribosomal RNA | 90, 10 |
| lN-hP, Nv | Chr | 568,401 | 4 bp→CGAA | noncoding (1397-1400/1489 nt) | ACNQK5_RS02750 ← | 16S ribosomal RNA | 100, 10 |
| lN-hP, Nv | Chr | 568,409 | G→A | noncoding (1392/1489 nt) | ACNQK5_RS02750 ← | 16S ribosomal RNA | 100, 10 |
| lN-hP, Nv | Chr | 568,412 | G→C | noncoding (1389/1489 nt) | ACNQK5_RS02750 ← | 16S ribosomal RNA | 100, 10 |
| lN-hP, Nv | Chr | 568,415 | 1 bp→CC | noncoding (1386/1489 nt) | ACNQK5_RS02750 ← | 16S ribosomal RNA | 100, 10 |
| lN-hP, Nv | Chr | 568,418 | Δ1 bp | noncoding (1383/1489 nt) | ACNQK5_RS02750 ← | 16S ribosomal RNA | 100, 10 |
| lN-hP, Nv | Chr | 568,423 | G→T | noncoding (1378/1489 nt) | ACNQK5_RS02750 ← | 16S ribosomal RNA | 100, 10 |
| lN-hP, Nv | Chr | 568,426 | C→T | noncoding (1375/1489 nt) | ACNQK5_RS02750 ← | 16S ribosomal RNA | 100, 10 |
| lN-hP, Nv | Chr | 568,428 | 2 bp→CA | noncoding (1372-1373/1489 nt) | ACNQK5_RS02750 ← | 16S ribosomal RNA | 100, 10 |
| lN-hP, Nv | Chr | 568,432 | C→A | noncoding (1369/1489 nt) | ACNQK5_RS02750 ← | 16S ribosomal RNA | 100, 10 |
| lN-hP, Nv | Chr | 568,434 | G→A | noncoding (1367/1489 nt) | ACNQK5_RS02750 ← | 16S ribosomal RNA | 100, 10 |
| lN-hP, Nv | Chr | 568,437 | T→C | noncoding (1364/1489 nt) | ACNQK5_RS02750 ← | 16S ribosomal RNA | 100, 10 |
| lN-hP, Nv | Chr | 568,477 | A→T | noncoding (1324/1489 nt) | ACNQK5_RS02750 ← | 16S ribosomal RNA | 100, 18 |
| lN-hP, Nv | Chr | 568,487 | C→A | noncoding (1314/1489 nt) | ACNQK5_RS02750 ← | 16S ribosomal RNA | 100, 18 |
| lN-hP, Nv | Chr | 568,489 | 1 bp→CA | noncoding (1312/1489 nt) | ACNQK5_RS02750 ← | 16S ribosomal RNA | 100, 18 |
| lN-hP, Nv | Chr | 568,497 | C→T | noncoding (1304/1489 nt) | ACNQK5_RS02750 ← | 16S ribosomal RNA | 100, 18 |
| lN-hP, Nv | Chr | 568,499 | G→T | noncoding (1302/1489 nt) | ACNQK5_RS02750 ← | 16S ribosomal RNA | 100, 18 |
| lN-hP, Nv | Chr | 568,518 | 2 bp→GA | noncoding (1282-1283/1489 nt) | ACNQK5_RS02750 ← | 16S ribosomal RNA | 100, 18 |
| lN-hP, Nv | Chr | 568,527 | C→G | noncoding (1274/1489 nt) | ACNQK5_RS02750 ← | 16S ribosomal RNA | 100, 18 |
| lN-hP, Nv | Chr | 568,530 | G→T | noncoding (1271/1489 nt) | ACNQK5_RS02750 ← | 16S ribosomal RNA | 100, 18 |

| | | | | | | | |
|---|---|---|---|---|---|---|---|
| lN-hP, Nv | Chr | 568,541 | C→A | noncoding (1260/1489 nt) | ACNQK5_RS02750 ← | 16S ribosomal RNA | 100, 18 |
| lN-hP, Nv | Chr | 568,544 | G→C | noncoding (1257/1489 nt) | ACNQK5_RS02750 ← | 16S ribosomal RNA | 100, 18 |
| lN-hP, Nv | Chr | 568,550 | T→C | noncoding (1251/1489 nt) | ACNQK5_RS02750 ← | 16S ribosomal RNA | 100, 18 |
| lN-hP, Nv | Chr | 568,552 | A→G | noncoding (1249/1489 nt) | ACNQK5_RS02750 ← | 16S ribosomal RNA | 100, 18 |
| lN-hP, Nv | Chr | 568,559 | 3 bp→ATA | noncoding (1240-1242/1489 nt) | ACNQK5_RS02750 ← | 16S ribosomal RNA | 100, 18 |
| lN-hP, Nv | Chr | 568,568 | Δ1 bp | noncoding (1233/1489 nt) | ACNQK5_RS02750 ← | 16S ribosomal RNA | 100, 18 |
| lN-hP, Nv | Chr | 568,577 | C→G | noncoding (1224/1489 nt) | ACNQK5_RS02750 ← | 16S ribosomal RNA | 100, 18 |
| lN-hP, Nv | Chr | 568,636 | A→T | noncoding (1165/1489 nt) | ACNQK5_RS02750 ← | 16S ribosomal RNA | 100, 6 |
| lN-hP, Nv | Chr | 2,023,374 | (AAAGGCA)15→19 | coding (118/243 nt) | ACNQK5_RS09745 → | hypothetical protein | 86.4, 144 |
| lN-hP, Nv | Chr | 2,367,866 | Δ144 bp | coding (3-146/213 nt) | ACNQK5_RS11275 ← | hypothetical protein | 100, 139 |
| lN-hP, Nv | Chr | 3,397,478 | C→A | noncoding (1271/1489 nt) | ACNQK5_RS16330 → | 16S ribosomal RNA | 100, 12 |
| lN-hP, Nv | Chr | 3,397,489 | G→T | noncoding (1282/1489 nt) | ACNQK5_RS16330 → | 16S ribosomal RNA | 100, 12 |
| lN-hP, Nv | Chr | 3,397,511 | G→A | noncoding (1304/1489 nt) | ACNQK5_RS16330 → | 16S ribosomal RNA | 100, 9 |
| lN-hP, Nv | Chr | 3,397,519 | A→G | noncoding (1312/1489 nt) | ACNQK5_RS16330 → | 16S ribosomal RNA | 100, 9 |
| lN-hP, Nv | Chr | 3,397,531 | T→A | noncoding (1324/1489 nt) | ACNQK5_RS16330 → | 16S ribosomal RNA | 100, 9 |
| lN-hP, Nv | Chr | 3,466,568 | C→A | A617A (GCG→GCT) | ACNQK5_RS16575 ← | beta strand repeat-containing protein | 85.7, 21 |
| lN-hP, Nv | Chr | 3,466,571 | G→A | D616D (GAC→GAT) | ACNQK5_RS16575 ← | beta strand repeat-containing protein | 89.5, 19 |
| lN-hP, Nv | P2 | 40,585 | A→G | pseudogene (241/732 nt) | PLSM73337038_RS00215 → | IS5-like element ISMae4 family transposase | 87.3, 118 |
| lN-hP, Nv | P2 | 40,793 | T→C | pseudogene (449/732 nt) | PLSM73337038_RS00215 → | IS5-like element ISMae4 family transposase | 84.0, 496 |
| hN-lP, Gv | Chr | 568,340 | G→A | noncoding (1461/1489 nt) | ACNQK5_RS02750 ← | 16S ribosomal RNA | 100, 10 |
| hN-lP, Gv | Chr | 568,374 | C→T | noncoding (1427/1489 nt) | ACNQK5_RS02750 ← | 16S ribosomal RNA | 100, 10 |
| hN-lP, Gv | Chr | 568,380 | C→A | noncoding (1421/1489 nt) | ACNQK5_RS02750 ← | 16S ribosomal RNA | 100, 10 |
| hN-lP, Gv | Chr | 568,387 | Δ1 bp | noncoding (1414/1489 nt) | ACNQK5_RS02750 ← | 16S ribosomal RNA | 100, 10 |
| hN-lP, Gv | Chr | 568,391 | +A | noncoding (1410/1489 nt) | ACNQK5_RS02750 ← | 16S ribosomal RNA | 100, 10 |
| hN-lP, Gv | Chr | 568,403 | 2 bp→AA | noncoding (1397-1398/1489 nt) | ACNQK5_RS02750 ← | 16S ribosomal RNA | 100, 10 |
| hN-lP, Gv | Chr | 568,409 | G→A | noncoding (1392/1489 nt) | ACNQK5_RS02750 ← | 16S ribosomal RNA | 100, 10 |
| hN-lP, Gv | Chr | 568,418 | Δ1 bp | noncoding (1383/1489 nt) | ACNQK5_RS02750 ← | 16S ribosomal RNA | 100, 10 |
| hN-lP, Gv | Chr | 568,423 | G→T | noncoding (1378/1489 nt) | ACNQK5_RS02750 ← | 16S ribosomal RNA | 100, 10 |
| hN-lP, Gv | Chr | 568,426 | C→T | noncoding (1375/1489 nt) | ACNQK5_RS02750 ← | 16S ribosomal RNA | 100, 10 |
| hN-lP, Gv | Chr | 568,429 | G→A | noncoding (1372/1489 nt) | ACNQK5_RS02750 ← | 16S ribosomal RNA | 100, 10 |
| hN-lP, Gv | Chr | 568,432 | C→A | noncoding (1369/1489 nt) | ACNQK5_RS02750 ← | 16S ribosomal RNA | 100, 10 |
| hN-lP, Gv | Chr | 568,434 | G→A | noncoding (1367/1489 nt) | ACNQK5_RS02750 ← | 16S ribosomal RNA | 100, 10 |
| hN-lP, Gv | Chr | 568,477 | A→T | noncoding (1324/1489 nt) | ACNQK5_RS02750 ← | 16S ribosomal RNA | 100, 13 |
| hN-lP, Gv | Chr | 568,487 | C→A | noncoding (1314/1489 nt) | ACNQK5_RS02750 ← | 16S ribosomal RNA | 100, 13 |
| hN-lP, Gv | Chr | 568,489 | 1 bp→CA | noncoding (1312/1489 nt) | ACNQK5_RS02750 ← | 16S ribosomal RNA | 100, 13 |

| | | | | | | | |
|---|---|---|---|---|---|---|---|
| hN-lP, Gv | Chr | 568,497 | C→T | noncoding (1304/1489 nt) | ACNQK5_RS02750 ← | 16S ribosomal RNA | 100, 13 |
| hN-lP, Gv | Chr | 568,499 | G→T | noncoding (1302/1489 nt) | ACNQK5_RS02750 ← | 16S ribosomal RNA | 100, 13 |
| hN-lP, Gv | Chr | 568,518 | 2 bp→GA | noncoding (1282-1283/1489 nt) | ACNQK5_RS02750 ← | 16S ribosomal RNA | 100, 13 |
| hN-lP, Gv | Chr | 568,527 | C→G | noncoding (1274/1489 nt) | ACNQK5_RS02750 ← | 16S ribosomal RNA | 100, 13 |
| hN-lP, Gv | Chr | 568,530 | G→T | noncoding (1271/1489 nt) | ACNQK5_RS02750 ← | 16S ribosomal RNA | 100, 13 |
| hN-lP, Gv | Chr | 568,541 | C→A | noncoding (1260/1489 nt) | ACNQK5_RS02750 ← | 16S ribosomal RNA | 100, 13 |
| hN-lP, Gv | Chr | 568,544 | G→C | noncoding (1257/1489 nt) | ACNQK5_RS02750 ← | 16S ribosomal RNA | 100, 13 |
| hN-lP, Gv | Chr | 568,550 | T→C | noncoding (1251/1489 nt) | ACNQK5_RS02750 ← | 16S ribosomal RNA | 100, 13 |
| hN-lP, Gv | Chr | 568,552 | A→G | noncoding (1249/1489 nt) | ACNQK5_RS02750 ← | 16S ribosomal RNA | 100, 13 |
| hN-lP, Gv | Chr | 568,559 | 3 bp→ATA | noncoding (1240-1242/1489 nt) | ACNQK5_RS02750 ← | 16S ribosomal RNA | 100, 13 |
| hN-lP, Gv | Chr | 568,568 | Δ1 bp | noncoding (1233/1489 nt) | ACNQK5_RS02750 ← | 16S ribosomal RNA | 100, 13 |
| hN-lP, Gv | Chr | 568,577 | C→G | noncoding (1224/1489 nt) | ACNQK5_RS02750 ← | 16S ribosomal RNA | 100, 12 |
| hN-lP, Gv | Chr | 2,023,374 | (AAAGGCA)15→19 | coding (118/243 nt) | ACNQK5_RS09745 → | hypothetical protein | 87.8, 355 |
| hN-lP, Gv | Chr | 2,367,866 | Δ144 bp | coding (3-146/213 nt) | ACNQK5_RS11275 ← | hypothetical protein | 100, 310 |
| *hN-lP, Gv* | *Chr* | *2,461,108* | *A→G* | *intergenic (-215/+53)* | *gvpC ← / ← gvpA* | *gas vesicle protein GvpC/gas vesicle structural protein GvpA* | *81.9, 205* |
| *hN-lP, Gv* | *Chr* | *2,461,116* | *2 bp→CA* | *intergenic (-223/+44)* | *gvpC ← / ← gvpA* | *gas vesicle protein GvpC/gas vesicle structural protein GvpA* | *80.1, 195* |
| hN-lP, Gv | Chr | 3,466,568 | C→A | A617A (GCG→GCT) | ACNQK5_RS16575 ← | beta strand repeat-containing protein | 98.4, 61 |
| hN-lP, Gv | Chr | 3,466,571 | G→A | D616D (GAC→GAT) | ACNQK5_RS16575 ← | beta strand repeat-containing protein | 98.3, 58 |
| hN-lP, Gv | P2 | 40,585 | A→G | pseudogene (241/732 nt) | PLSM73337038_RS00215 → | IS5-like element ISMae4 family transposase | 84.8, 283 |
| hN-lP, Gv | P2 | 40,793 | T→C | pseudogene (449/732 nt) | PLSM73337038_RS00215 → | IS5-like element ISMae4 family transposase | 86.5, 1137 |
| hN-lP, Nv | Chr | 2,367,866 | Δ144 bp | coding (3-146/213 nt) | ACNQK5_RS11275 ← | hypothetical protein | 100, 348 |
| *hN-lP, Nv* | *Chr* | *2,462,489* | *C→T* | *M1M (ATG→ATA) †* | *gvpA ←* | *gas vesicle structural protein GvpA* | *100, 384* |
| hN-lP, Nv | Chr | 4,502,481 | (TTTGCCT)9→10 | intergenic (+168/+59) | ACNQK5_RS21635 → / ← ACNQK5_RS21640 | four helix bundle protein/type ISP restriction/modification enzyme | 92.5, 387 |
| hN-lP, Nv | P1 | 1 | Δ18,380 bp | | [PLSM72508198_RS00005]–[PLSM72508198_RS00005] | 32 genes | N.a. |
| hN-lP, Nv | P2 | 40,585 | A→G | pseudogene (241/732 nt) | PLSM73337038_RS00215 → | IS5-like element ISMae4 family transposase | 88.6, 255 |
| hN-lP, Nv | P2 | 40,793 | T→C | pseudogene (449/732 nt) | PLSM73337038_RS00215 → | IS5-like element ISMae4 family transposase | 91.7, 990 |
| lN-lP, Gv | Chr | 2,023,374 | (AAAGGCA)15→19 | coding (118/243 nt) | ACNQK5_RS09745 → | hypothetical protein | 85.4, 377 |
| lN-lP, Gv | Chr | 2,367,866 | Δ144 bp | coding (3-146/213 nt) | ACNQK5_RS11275 ← | hypothetical protein | 100, 421 |

| | | | | | | | |
|---|---|---|---|---|---|---|---|
| *lN-lP, Gv* | *Chr* | *2,462,329* | *G→T* | *L55I (CTC→ATC)* | *gvpA ←* | *gas vesicle structural protein GvpA* | *99.8, 475* |
| *lN-lP, Gv* | *Chr* | *2,462,342* | *A→C* | *S50S (TCT→TCG)* | *gvpA ←* | *gas vesicle structural protein GvpA* | *99.6, 473* |
| **lN-lP, Gv** | **Chr** | **3,210,317** | **(C)12→11** | **intergenic (-199/-259)** | **ACNQK5_RS15405 ← / → ACNQK5_RS15410** | **PEP-CTERM sorting domain-containing protein/Uma2 family endonuclease** | **94.0, 152** |
| lN-lP, Gv | Chr | 3,466,568 | C→A | A617A (GCG→GCT) | ACNQK5_RS16575 ← | beta strand repeat-containing protein | 83.7, 43 |
| lN-lP, Gv | Chr | 3,466,571 | G→A | D616D (GAC→GAT) | ACNQK5_RS16575 ← | beta strand repeat-containing protein | 92.7, 41 |
| **lN-lP, Gv** | **Chr** | **3,802,561** | **G→T** | **G554V (GGT→GTT)** | **ACNQK5_RS18290 →** | **peptidase domain-containing ABC transporter** | **99.8, 408** |
| **lN-lP, Gv** | **Chr** | **4,213,818** | **G→C** | **L71V (CTT→GTT)** | **purF ←** | **amidophosphoribosyltransferase** | **100, 485** |
| lN-lP, Gv | P1 | 3,665 | G→A | R128Q (CGA→CAA)<br>D7N (GAT→AAT) | PLSM72508198_RS00035 →<br>PLSM72508198_RS00040 → | hypothetical protein<br>hypothetical protein | 87.5, 16 |
| lN-lP, Gv | P1 | 3,706 | G→A | K20K (AAG→AAA) | PLSM72508198_RS00040 → | hypothetical protein | 87.5, 16 |
| lN-lP, Gv | P1 | 3,709 | A→T | P21P (CCA→CCT) | PLSM72508198_RS00040 → | hypothetical protein | 87.5, 16 |
| lN-lP, Gv | P1 | 3,715 | C→T | Y23Y (TAC→TAT) | PLSM72508198_RS00040 → | hypothetical protein | 87.5, 16 |
| lN-lP, Gv | P1 | 3,717 | 3 bp→GTG | coding (71-73/168 nt) | PLSM72508198_RS00040 → | hypothetical protein | 87.5, 16 |
| lN-lP, Gv | P2 | 40,585 | A→G | pseudogene (241/732 nt) | PLSM73337038_RS00215 → | IS5-like element ISMae4 family transposase | 90.1, 503 |
| lN-lP, Gv | P2 | 40,793 | T→C | pseudogene (449/732 nt) | PLSM73337038_RS00215 → | IS5-like element ISMae4 family transposase | 91.2, 2209 |
| lN-lP, Nv | Chr | 568,340 | G→A | noncoding (1461/1489 nt) | ACNQK5_RS02750 ← | 16S ribosomal RNA | 100, 11 |
| lN-lP, Nv | Chr | 568,380 | C→A | noncoding (1421/1489 nt) | ACNQK5_RS02750 ← | 16S ribosomal RNA | 100, 11 |
| lN-lP, Nv | Chr | 568,382 | G→A | noncoding (1419/1489 nt) | ACNQK5_RS02750 ← | 16S ribosomal RNA | 100, 11 |
| lN-lP, Nv | Chr | 568,403 | 2 bp→AA | noncoding (1397-1398/1489 nt) | ACNQK5_RS02750 ← | 16S ribosomal RNA | 100, 11 |
| lN-lP, Nv | Chr | 568,409 | G→A | noncoding (1392/1489 nt) | ACNQK5_RS02750 ← | 16S ribosomal RNA | 100, 11 |
| lN-lP, Nv | Chr | 568,418 | Δ1 bp | noncoding (1383/1489 nt) | ACNQK5_RS02750 ← | 16S ribosomal RNA | 100, 11 |
| lN-lP, Nv | Chr | 568,429 | G→A | noncoding (1372/1489 nt) | ACNQK5_RS02750 ← | 16S ribosomal RNA | 100, 11 |
| lN-lP, Nv | Chr | 568,434 | G→A | noncoding (1367/1489 nt) | ACNQK5_RS02750 ← | 16S ribosomal RNA | 100, 11 |
| lN-lP, Nv | Chr | 568,477 | A→T | noncoding (1324/1489 nt) | ACNQK5_RS02750 ← | 16S ribosomal RNA | 100, 12 |
| lN-lP, Nv | Chr | 568,487 | C→A | noncoding (1314/1489 nt) | ACNQK5_RS02750 ← | 16S ribosomal RNA | 100, 12 |
| lN-lP, Nv | Chr | 568,489 | 1 bp→CA | noncoding (1312/1489 nt) | ACNQK5_RS02750 ← | 16S ribosomal RNA | 100, 12 |
| lN-lP, Nv | Chr | 568,497 | C→T | noncoding (1304/1489 nt) | ACNQK5_RS02750 ← | 16S ribosomal RNA | 100, 12 |
| lN-lP, Nv | Chr | 568,499 | G→T | noncoding (1302/1489 nt) | ACNQK5_RS02750 ← | 16S ribosomal RNA | 100, 12 |
| lN-lP, Nv | Chr | 568,518 | 2 bp→GA | noncoding (1282-1283/1489 nt) | ACNQK5_RS02750 ← | 16S ribosomal RNA | 100, 12 |
| lN-lP, Nv | Chr | 568,527 | C→G | noncoding (1274/1489 nt) | ACNQK5_RS02750 ← | 16S ribosomal RNA | 100, 12 |
| lN-lP, Nv | Chr | 568,530 | G→T | noncoding (1271/1489 nt) | ACNQK5_RS02750 ← | 16S ribosomal RNA | 100, 12 |
| lN-lP, Nv | Chr | 568,541 | C→A | noncoding (1260/1489 nt) | ACNQK5_RS02750 ← | 16S ribosomal RNA | 100, 12 |
| lN-lP, Nv | Chr | 568,544 | G→C | noncoding (1257/1489 nt) | ACNQK5_RS02750 ← | 16S ribosomal RNA | 100, 12 |
| lN-lP, Nv | Chr | 568,550 | T→C | noncoding (1251/1489 nt) | ACNQK5_RS02750 ← | 16S ribosomal RNA | 100, 12 |

| | | | | | | | |
|---|---|---|---|---|---|---|---|
| lN-lP, Nv | Chr | 568,552 | A→G | noncoding (1249/1489 nt) | ACNQK5_RS02750 ← | 16S ribosomal RNA | 100, 12 |
| lN-lP, Nv | Chr | 568,560 | 2 bp→TA | noncoding (1240-1241/1489 nt) | ACNQK5_RS02750 ← | 16S ribosomal RNA | 100, 12 |
| lN-lP, Nv | Chr | 568,568 | Δ1 bp | noncoding (1233/1489 nt) | ACNQK5_RS02750 ← | 16S ribosomal RNA | 100, 12 |
| lN-lP, Nv | Chr | 2,023,374 | (AAAGGCA)15→19 | coding (118/243 nt) | ACNQK5_RS09745 → | hypothetical protein | 84.2, 238 |
| lN-lP, Nv | Chr | 2,367,866 | Δ144 bp | coding (3-146/213 nt) | ACNQK5_RS11275 ← | hypothetical protein | 100, 270 |
| lN-lP, Nv | Chr | 3,466,568 | C→A | A617A (GCG→GCT) | ACNQK5_RS16575 ← | beta strand repeat-containing protein | 94.6, 37 |
| lN-lP, Nv | Chr | 3,466,571 | G→A | D616D (GAC→GAT) | ACNQK5_RS16575 ← | beta strand repeat-containing protein | 91.4, 35 |
| lN-lP, Nv | P2 | 40,585 | A→G | pseudogene (241/732 nt) | PLSM73337038_RS00215 → | IS5-like element ISMae4 family transposase | 86.8, 197 |
| lN-lP, Nv | P2 | 40,793 | T→C | pseudogene (449/732 nt) | PLSM73337038_RS00215 → | IS5-like element ISMae4 family transposase | 90.1, 710 |

[a]Nucleotide substitutions (‘Mutation column’) are reported relative to the reference genome sequence, and codon changes (‘Description’ column) are shown in coding-strand orientation.

[b]Values before the comma indicate the percentage of reads supporting the mutation, and values after the comma indicate the total read count at that position, as reported by *breseq*.

Abbreviations: hN-hP, high N & high P; lN-hP, low N & high P; hN-lP, high N & low P; lN-lP, low N & low P; Gv, gas-vacuolate NIES-298; Nv, non-vacuolate NIES-298; Chr, chromosome; P1, plasmid 1; P2, plasmid 2; N.a., not applicable.
